# Nephron progenitor maintenance is controlled through FGFs and Sprouty1 Interaction

**DOI:** 10.1101/2020.03.26.010421

**Authors:** Sung-Ho Huh, Ligyeom Ha, Hee-Seong Jang

**Affiliations:** Department of Neurological Sciences, University of Nebraska Medical Center, Omaha, NE, 68198, USA; Cellular & Integrative Physiology, University of Nebraska Medical Center, Omaha, NE, 68198, USA; Holland Regenerative Medicine Program, University of Nebraska Medical Center, Omaha, NE, 68198, USA

**Keywords:** FGF signaling, Spry1, nephron progenitor cells, development

## Abstract

The nephron progenitor cells (NPCs) give rise to all segments of functional nephrons and are of great interest due to their potential as a source for novel treatment strategies for kidney disease. Fibroblast growth factor (FGF) signal plays pivotal roles in generating and maintaining NPCs during kidney development. However, molecule(s) regulating FGF signal during nephron development is not known. Sprouty (SPRY) is an antagonist of receptor tyrosine kinases. During kidney development, SPRY1 is expressed in the ureteric buds (UBs) and regulates UB branching by antagonizing Ret-GDNF signal. Here, we provide evidence that SPRY1 expressed in NPCs modulates activity of FGF signal in NPCs and regulates NPC stemness. Haploinsufficiency of *Spry1* rescues bilateral renal agenesis and premature NPC differentiation caused by loss of *Fgf9* and *Fgf20*. In addition, haploinsufficiency of *Spry1* rescues NPC proliferation and cell death defects induced by loss of *Fgf9* and *Fgf20*. In the absence of SPRY1, FGF9 and FGF20, another FGF ligand, FGF8 promotes nephrogenesis. Deleting both *Fgf8* and *Fgf20* results in kidney agenesis and defects in NPC proliferation and cell death. Rescue of loss of Fgf9 and Fgf20 induced renal agenesis by *Spry1* haploinsufficiency was reversed when one copy of *Fgf8* was deleted. These findings indicate the importance of the balance between positive and negative signal during NPC maintenance. Failure of the balance may underlie some human congenital kidney malformation.

**Significance Statement:** Nephrons are functional units of kidney to filter blood to excrete wastes and regulate osmolarity and ion concentrations. Nephrons are derived from nephron progenitors. Nephron progenitors are depleted during kidney development which makes it unable to regenerate nephrons. Therefore, understanding signaling molecules regulating nephron progenitor generation and maintenance is of great interest for the future kidney regenerative medicine. Here, we show that Sprouty1 regulates nephron progenitor maintenance by inhibiting FGF signal. Deletion of Sprouty1 rescues renal agenesis and nephron progenitor depletion in the Fgf9/20 loss-of-function kidneys. Further decrease of FGF signal by deleting one copy of Fgf8 makes kidneys irresponsive to Sprouty1 resulting in failure of nephron progenitor maintenance. This study thus identifies the reciprocal function of FGF-Sprouty1 signal during nephron progenitor development.

## Introduction

Receptor tyrosine kinases (RTK) are activated upon binding of their cognate ligands and regulate many aspects of organogenesis. The intracellular signals initiated by RTK activation determine cellular behavior such as proliferation, survival, cell fate determination and morphogenesis. Mis-regulation of RTK activity leads to the onset and progression of a wide-range of disease such as diabetes, inflammation, bone disorders, atherosclerosis, angiogenesis and various cancers (1, 2). Therefore, tight regulation of the activity of RTK should be guaranteed during development and homeostasis (3).

RTK plays crucial role during kidney development. glial cell line-derived neurotrophic factor (GDNF) produced in the nephron progenitor cells (NPCs) activates RET/GFRα1 RTK/co-receptor in the ureteric bud (UB) tips (4). Deletion of *Gdnf, Ret*, or *Gfra1* during development results in renal agenesis due to failure of UB induction (5–8). UB branching is sensitive to the level of GDNF since haploinsufficiency of GDNF or decreasing expression of GDNF due to loss of *Fras1* produces unilateral renal agenesis or hypoplasia (9, 10). On the other hand, FGF9 and FGF20 activate FGFR1 and FGFR2 in NPCs and maintain the stemness of NPCs. Loss of FGF20 in human, loss of *Fgf9* and *Fgf20* in mice, or metanephric mesenchyme specific deletion of *Fgfr1* and *Fgfr2* causes bilateral renal agenesis (11–13). In addition, reduction in *Fgf9* and *Fgf20* levels results in loss of NPCs and premature differentiation indicating that stemness of NPCs are sensitive to the copy number of *Fgf* genes (12).

Sprouty (SPRY) is a negative feedback regulator of RTK signaling. During kidney development, SPRY negatively regulates activity of GDNF-RET signal. Either knock-out or antisense oligonucleotides treatment of *Spry1* generates supernumerary and dilated UBs (14, 15). In addition, ectopic expression of SPRY2 in the developing kidney leads to renal hypoplasia and agenesis (16). Furthermore, deleting *Spry1* restores kidneys in the *Gdnf* (or its expression level) or *Ret* knock-out mice (17–19). When both GDNF/RET and SPRY signal is depleted, FGF10 promotes UB branching (18). Although these studies point to the importance of SPRY1 in the UB branching, the function of SPRY1 in the NPCs *in vivo* remains to be determined.

Here, we show that SPRY1 regulates NPC maintenance by modulating FGF signal. Both germ line and NPC specific deletion of *Spry1* rescue renal agenesis in *Fgf9*^-/-^;*Fgf20*^-/-^ embryos. We also show that FGF8 is another FGF ligand promoting NPC maintenance. Combinatory gene deletion analyses show that balance of FGFs and Spry1 level is critical to maintain NPCs.

## Results

### Loss of *Spry1* partially restores kidney phenotypes caused by loss of *Fgf9* and *Fgf20*

To test whether *Spry1* regulates *Fgf9* and *Fgf20* activity during kidney development, we generated *Fgf9, Fgf20*, and *Spry1* triple compound mutants. In this study, we used *Fgf20*^-/-^ embryos as controls. Previously, we identified that *Fgf20*^-/-^ embryos produce 15% smaller kidneys than *Fgf20*^-/+^ with decreased number of nephrons (12). Kidney size of *Fgf20*^-/-^ and *Fgf20*^-/-^;*Spry1*^-/+^ embryos were comparable at E18.5 (100.3±8.0% vs. 101.3±9.7%, n=12, ns, Figs. 1A, B, Q, R). Histological morphology appears normal in both genotypes (Figs. 1G and H). In *Fgf9*^-/+^;*Fgf20*^-/-^ embryos, 10 out of 18 kidneys were hypoplastic and 1 kidney was aplastic and kidney size was decreased to 61.8±30.1% compared to *Fgf20*^-/-^ (p<0.001) (Figs. 1C, Q, R). *Fgf9*^-/+^;*Fgf20*^-/-^ kidneys were also cystic (Fig. 1I). In *Fgf9*^-/+^;*Fgf20*^-/-^;*Spry1*^-/+^ embryos, 5 out of 18 kidneys were hypoplastic and kidney size was significantly restored compared to *Fgf9*^-/+^;*Fgf20*^-/-^ embryos (87.4±15.0%, p<0.005 compared to *Fgf9*^-/+^;*Fgf20*^-/-^ Figs. 1D, Q, R). Histologically, *Fgf9*^-/+^:*Fgf20*^-/-^;*Spry1*^-/+^ kidneys were comparable to controls (Fig. 1J). However, kidney size was not restored back to control size (p<0.05 compared to *Fgf20*^-/-^)(Fig. 1R). All kidneys in *Fgf9*^-/-^;*Fgf20*^-/-^ embryos were missing (n=8) (Figs. 1E and Q). Interestingly, all *Fgf9*^-/-^;*Fgf20*^-/-^;*Spry1*^-/+^ embryos generated kidneys. Among 14 kidneys analyzed, 4 kidneys were normal and 8 kidneys were hypoplastic (Figs. 1F and Q). Size of the kidneys in *Fgf9*^-/-^;*Fgf20*^-/-^;*Spry1*^-/+^ embryos was 64±18.9% compared to controls, which is significantly increased compared to in *Fgf9*^-/-^;*Fgf20*^-/-^ embryos (p<0.005) and comparable to in *Fgf9*^-/+^;*Fgf20*^-/-^ embryos (p>0.75) (Fig. 1R). Histological analysis showed that kidney structures were comparable to controls (Fig. 1K). Nephron progenitors were repopulated in *Fgf9*^-/+^;*Fgf20*^-/-^;*Spry1*^-/+^ and *Fgf9*^-/-^;*Fgf20*^-/-^;*Spry1*^-/+^ kidneys compared to *Fgf9*^-/+^;*Fgf20*^-/-^;*Spry1*^-/+^ and *Fgf9*^-/-^;*Fgf20*^-/-^;*Spry1*^-/+^, respectively (Figs. 1L-P’). These data indicate that deleting one copy of *Spry1* partially restores nephrogenesis caused by loss of *Fgf9* and *Fgf20*.

**Fig. 1.**
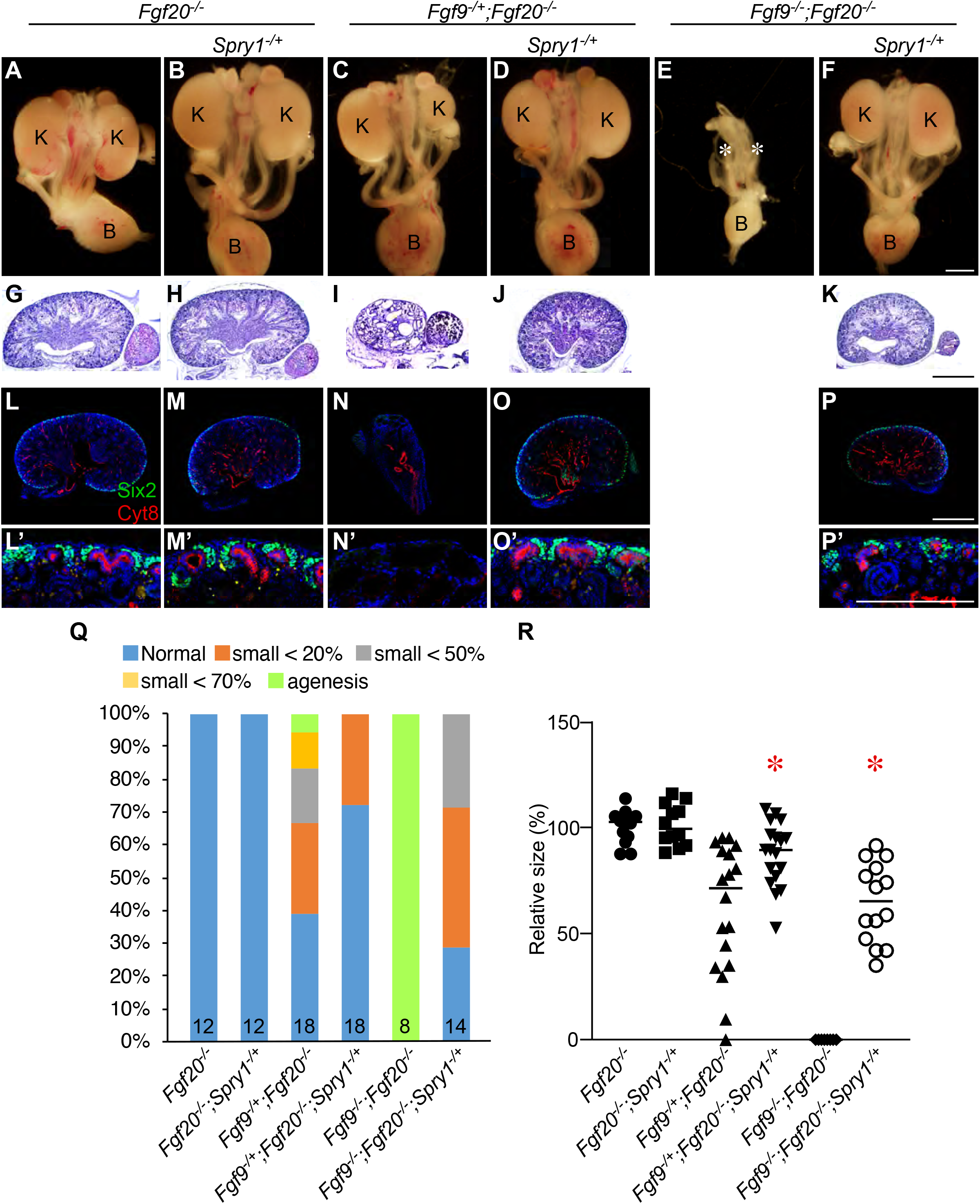
Haploinsufficiency of *Spry1* rescues renal phenotypes caused by loss of *Fgf9* and *Fgf20*. (**A-F**) Morphology of urogenital system in E18.5 *Fgf20*^-/-^ (**A**), *Fgf20*^-/-^;*Srpy1*^-/+^ (**B**), *Fgf9*^-/+^;*Fgf20*^-/-^ (**C**), *Fgf9*^-/+^;*Fgf20*^-/-^;*Spry1*^-/+^ (**D**), *Fgf9*^-/-^;*Fgf20*^-/-^ (**E**), and *Fgf9*^-/-^;*Fgf20*^-/-^;*Spry1*^-/+^ (**F**). (G-K) Hematoxylin & Eosin staining of *Fgf20*^-/-^ (**G**), *Fgf20*^-/-^;*Srpy1*^-/+^ (**H**), *Fgf9*^-/+^;*Fgf20*^-/-^ (**I**), *Fgf9*^-/+^;*Fgf20*^-/-^;*Spry1*^-/+^ (**J**), and *Fgf9*^-/-^;*Fgf20*^-/-^;*Spry1*^-/+^ (**K**) kidneys. (**L-P**) Sections of E18.5 kidneys stained with Cytokeratin 8 (Cyt8) and Six2 in *Fgf20*^-/-^ (**L**), *Fgf20*^-/-^;*Srpy1*^-/+^ (**M**), *Fgf9*^-/+^;*Fgf20*^-/-^ (**N**), *Fgf9*^-/+^;*Fgf20*^-/-^;*Spry1*^-/+^ (**O**), and *Fgf9*^-/-^;*Fgf20*^-/-^;*Spry1*^-/+^ (**P**) kidneys. (**L’-P’**) high power image of nephrogenic zone from *Fgf20*^-/-^ (**L’**), *Fgf20*^-/-^;*Srpy1*^-/+^ (**M’**), *Fgf9*^-/+^;*Fgf20*^-/-^ (**N’**), *Fgf9*^-/+^;*Fgf20*^-/-^;*Spry1*^-/+^ (**O’**), and *Fgf9*^-/-^;*Fgf20*^-/-^;*Spry1*^-/+^ (**P’**) kidneys. (**Q**) percentage of kidney phenotypes. (**R**) Relative kidney size of *Fgf20*^-/-^ (n=12), *Fgf20*^-/-^;*Srpy1*^-/+^ (n=12), *Fgf9*^-/+^;*Fgf20*^-/-^ (n=18), *Fgf9*^-/+^;*Fgf20*^-/-^;Spry1^-/+^ (n=18), *Fgf9*^-/-^;*Fgf20*^-/-^ (n=8), and *Fgf9*^-/-^;Fgf20-^/-^;*Spry1*^-/+^ (n=14). K, kidney, B, bladder. * in E indicates loss of kidneys. *P < 0.01. Scale bar, 100μm.

*Spry1* is reported to be expressed in both UBs and NPCs during kidney development (14, 15). We observed that *Spry1* was expressed in both UBs and NPCs at E11.5 and its expression was decreased in NPCs at E13.5 and E14.5 kidneys (Fig. S1). *Spry1* expressed in the UBs negatively regulates GDNF-RET induced UB branch morphogenesis (17, 18). We hypothesized that *Spry1* expressed in the NPCs antagonizes FGF9 and FGF20 induced NPC maintenance. To investigate this, we conditionally deleted *Spry1* in NPCs using *Fgf20^Cre^* knock-in mouse line, which serves both loss of *Fgf20* and Cre driver of NPC (12). Cre recombinase in *Fgf20^Cre^* was active only in NPCs and their derivatives (Fig. S2).

Kidney size of *Fgf20^Cre/-^* and *Fgf20^Cre/-^;Spry1^fl/+^* embryos were comparable at E18.5 (99.7±14.0% vs. 99.0±10.5%, n=12, ns, Figs. 2A, B, G, H). In *Fgf9*^-/+^;*Fgf20^Cre/-^* embryos, 3 out of 16 kidneys were hypoplastic and 2 kidney was aplastic and kidney size was decreased to 75.7±32.1% compared to *Fgf20^Cre/-^* (p<0.05) (Figs. 2C, G, H). In *Fgf9^-/+^;Fgf20^Cre/-^;Spry1^fl/+^* embryos, 2 out of 18 kidneys were hypoplastic and kidney size was 87.6±22.9% compared to *Fgf20^Cre/-^* (p>0.22) (Figs. 2D, G, H). All kidneys in *Fgf9*^-/-^;*Fgf20^Cre/-^* embryos were missing (n=12) (Figs. 2E and G). 6 out of 8 kidneys in *Fgf9^-/-^;Fgf20^Cre/-^;Spry1^fl/+^* embryos generated kidneys. Among 8 kidneys analyzed, 2 kidneys were normal, 4 kidneys were hypoplastic, and 2 kidneys were missing (Figs. 2F and G). Size of the kidneys in *Fgf9*^-/-^;*Fgf20^cre/-^;Spry1^fl/+^* embryos was 56.2±41.6% compared to *Fgf20^Cre/-^*, which is significantly increased compared to in *Fgf9*^-/-^;*Fgf20*^-/-^ embryos (p<0.005) (Fig 2H). Together, these data indicate that phenotypic rescue of NPC specific *Spry1* deletion recapitulates whole body deletion of *Spry1* suggesting SPRY1 in NPCs antagonizes activity of FGF9 and FGF20 to regulate kidney development.

**Fig. 2.**
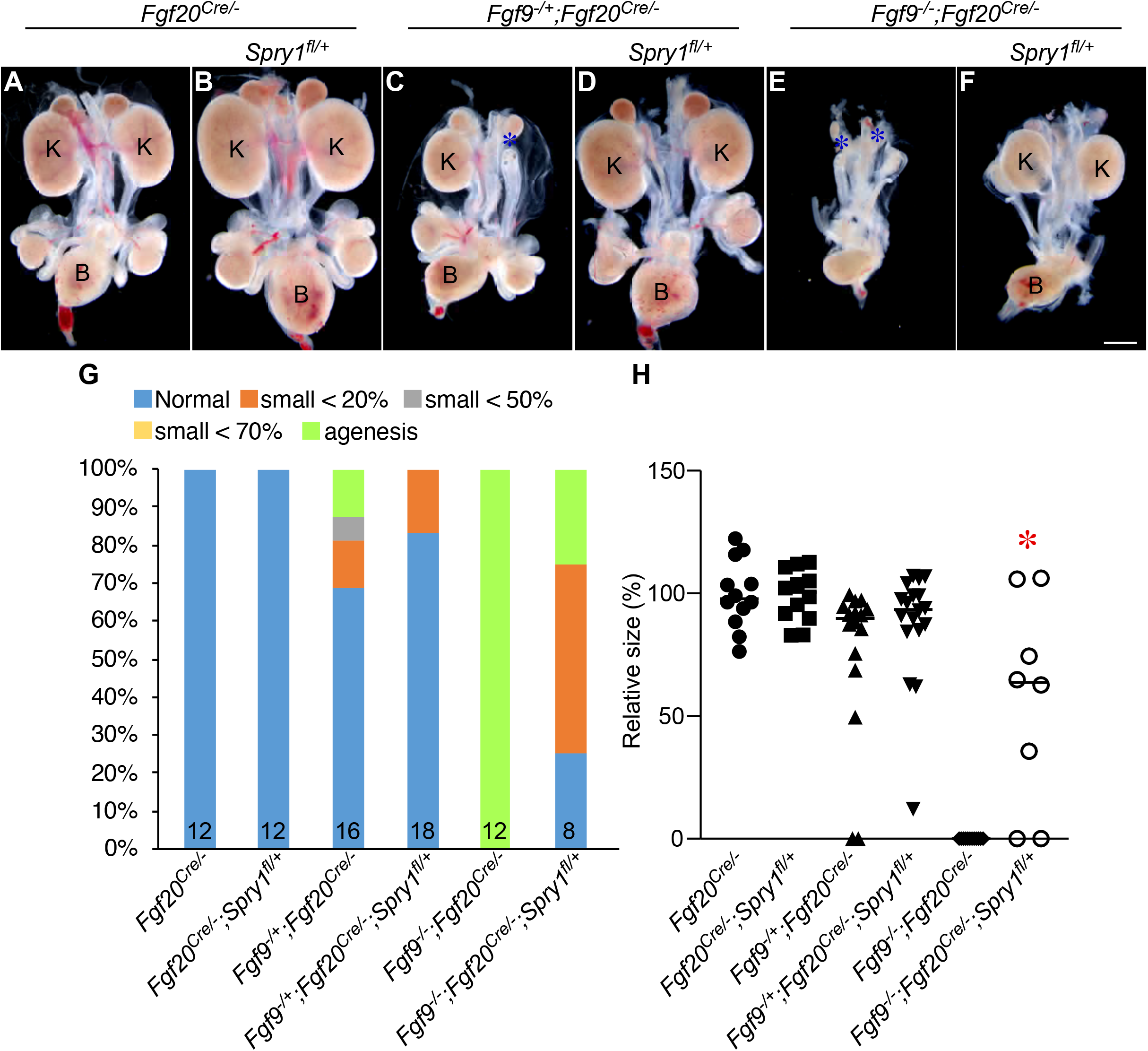
*Spry1* in nephron progenitor cells antagonizes *Fgf9* and *Fgf20* induced nephrogenesis. (**A-F**) Morphology of urogenital system in E18.5 *Fgf20^Cre/-^* (**A**), *Fgf20^Cre/-^;Srpy1^fl/+^* (**B**), *Fgf9*^-/+^;*Fgf20^Cre/-^*. (**Q**) percentage of kidney phenotypes. (**R**) Relative kidney size of *Fgf20^Cre/-^* (n=12), *Fgf20^Cre/-^;Srpy1^fl/+^* (n=12), *Fgf9*^-/+^;*Fgf20^Cre/-^* (n=16), *Fgf9*^-/+^;*Fgf20^Cre/-^;Spry1^fl/+^* (n=18), *Fgf9*^-/-^;*Fgf20^Cre/-^* (n=12), and *Fgf9*^-/-^;*Fgf20^Cre/-^;Spry1^fl/+^* (n=8). K, kidney, B, bladder. * in E indicates loss of kidneys. *P < 0.01. Scale bar: 100μm.

### FGF9 and FGF20 are required for nephron progenitor survival and proliferation

Previously, we identified that loss of *Fgf9* and *Fgf20* resulted in NPC cell death (12). To further analyze their roles during kidney development, we performed proliferation and cell death analyses of NPCs at the time of NPC induction. At E10.5, number (67.4±15.3, 54.2±3.9, 61.5±11.2, 58.8±9.8 in *Fgf9*^-/+^;*Fgf20*^-/+^, *Fgf9*^-/-^;*Fgf20*^-/+^, *Fgf9*^-/-^;Fgf20^-/-^, *Fgf9*^-/-^;*Fgf20*^-/-^, respectively, n=4, ns), proliferation rate (22.5±7.4, 18.5±4.4, 20.5±4.5, 17.1±1.1, n=4, ns), and cell death rate (0.0±0.0, 0.0±0.0, 0.2±0.2, 0.5±0.6, n=4, ns) of Six2+ NPCs were comparable to all genotypes (Fig. S3). At E11.5, number of NPCs were comparable in *Fgf9*^-/+^;*Fgf20*^-/+^, *Fgf9*^-/-^;*Fgf20*^-/+^, *Fgf9*^+/-^;*Fgf20*^-/-^ embryos (70.9±8.9, 67.3±7.2, 74.4±2.5, respectively, n=3, ns)(Figs. S4A-C and M). However, number of NPCs in *Fgf9*^-/-^;*Fgf20*^-/-^ embryos (38.4±7.6, n=3, p<0.01) was significantly decreased compared to *Fgf9*^-/+^;*Fgf20*^-/+^ embryos (Figs. S4D and M). Proliferation rate of NPCs in *Fgf9*^-/-^;*Fgf20*^-/-^ embryos (19.5±1.6, n=3, p<0.01) was also significantly decreased compared to *Fgf9*^-/+^;*Fgf20*^-/+^ embryos (27.9±2.2, n=3) (Figs. S4H and N). Proliferation rates of NPCs in *Fgf9*^-/-^;*Fgf20*^-/+^ (28.6±1.9, n=3) and *Fgf9*^-/+^;*Fgf20*^-/-^ (27.9±1.4, n=3) embryos were comparable to *Fgf9*^-/+^;*Fgf20*^-/+^ embryos (Figs. 4E-G and N). Cell death rates of NPCs in *Fgf9*^-/+^;*Fgf20*^-/+^ and *Fgf9*^-/-^;*Fgf20*^-/+^ embryos were comparable (0.3±0.3, 0.7±0.6, n=3, respectively, ns) (Figs. S4I, J, O). Cell death rate of NPCs in *Fgf9*^-/+^;*Fgf20*^-/-^ (2.4±0.8, n=3 p<0.05) embryos was increased compared to *Fgf9*^-/+^;*Fgf20*^-/+^ embryos (Figs. S4K and O). Cell death rate of NPCs was further increased in *Fgf9*^-/-^;*Fgf20*^-/-^ (8.5±3.0, n=3), p<0.01) embryos compared to *Fgf9*^-/+^;*Fgf20*^-/+^ (p<0.01) and *Fgf9*^-/+^;*Fgf20*^-/-^ (p<0.05) embryos (Figs. S4L and O).

### *Fgf9* and *Fgf20* regulate ureteric bud branching

We also investigated whether FGF9 and FGF20 regulate genes required for UB branching. At E10.5, *Ret* was expressed in UBs and its expression was comparable to all the genotypes (Figs. S5A-D). At E11.5, *Ret+* UBs were bifurcated in *Fgf9*^-/+^;*Fgf20*^-/+^ and *Fgf9*^-/-^;*Fgf20*^-/+^ embryos (Figs. S5E and F). However, in *Fgf9*^-/+^;*Fgf20*^-/-^ embryos, UBs were not yet bifurcated (Fig. S5G) and in *Fgf9*^-/-^;*Fgf20*^-/-^ embryos, UBs started to regress (Fig. S5H). *Gdnf* was highly expressed in NPCs of *Fgf9*^-/+^;*Fgf20*^-/+^ and *Fgf9^-/-;^Fgf20^-/+^ embryos (Figs. SI and J). However, expression of *Gdnf* was decreased in *Fgf9*^-/+^;*Fgf20*^-/-^* embryos (Fig. S5K) and diminished in *Fgf9*^-/-^;*Fgf20*^-/-^ embryos (Fig. S5L). *Etv4* and *Etv5* are transcription factors regulated by GDNF/RET during UB branching and responsible for UB branching (18, 20–22). Therefore, we tested whether decrease of *Gdnf* affect expression of *Etv4* and *Etv5*. At E11.5, expression of *Etv4* and *Etv5* were significantly decreased in *Fgf9*^-/+^;*Fgf20*^-/-^ embryos and diminished in *Fgf9*^-/-^;*Fgf20*^-/-^ embryos (Figs. S5M-S). Together, these data indicate that *Fgf9* and *Fgf20* are required for UB branching.

### Reduction of *Spry1* level restores nephron progenitor proliferation and survival in *Fgf9* and *Fgf20* double mutant kidneys

Next, we investigated whether deleting one copy of *Spry1* rescues NPC number, proliferation and cell death defects caused by loss of *Fgf9* and *Fgf20*. Number of NPCs were comparable in both *Fgf20*^-/-^ and *Fgf20*^-/-^;*Spry1*^-/+^ embryos (63.3±5.7, 60.8±9.4, n=3, respectively, ns) (Figs. 3A, D, G). Number of NPCs in *Fgf9*^-/+^;*Fgf20*^-/-^;*Spry1*^-/+^ embryos was increased compared to that of *Fgf9*^-/+^;*Fgf20*^-/-^ embryos (44.6±12.0, 60.0±4.8, n=5, respectively, p<0.05) (Figs. 3C, D, G). Also, Number of NPCs in *Fgf9*^-/-^;*Fgf20*^-/-^;*Spry1*^-/+^ embryos was increased compared to that of *Fgf9*^-/-^;*Fgf20*^-/-^ embryos (24.9±8.6, 54.0±2.0, n=5, n=4 respectively, p<0.001) (Figs. 3E, F, G). Of note, number of NPCs in *Fgf9*^-/+^;*Fgf20*^-/-^;*Spry1*^-/+^ and *Fgf9*^-/-^;*Fgf20*^-/-^;*Spry1*^-/+^ embryos was restored close to that of *Fgf20*^-/-^ embryos (p>0.5). Proliferation indexes were comparable to both *Fgf20*^-/-^ and *Fgf20*^-/-^;*Spry1*^-/+^ embryos (21.9±4.7, 23.2±4.0, n=3, respectively, ns) (Figs. 3H, I, N). Proliferation index in *Fgf9*^-/+^;*Fgf20*^-/-^;*Spry1*^-/+^ embryos was increased compared to that of *Fgf9*^-/+^;*Fgf20*^-/-^ embryos (13.7±4.6, 21.4±3.8, n=5, respectively, p<0.05) (Figs. 3J, K, N). Also, proliferation index in *Fgf9*^-/-^;*Fgf20*^-/-^;*Spry1*^-/+^ embryos was increased compared to that of *Fgf9*^-/-^;*Fgf20*^-/-^ embryos (8.7±4.1, 21.0±1.2, n=5, n=4 respectively, p<0.001) (Fig. 3 L, M, N). Of note, Proliferation indexes in *Fgf9*^-/+^;*Fgf20*^-/-^;*Spry1*^-/+^ and *Fgf9*^-/-^;*Fgf20*^-/-^;*Spry1*^-/+^ embryos was restored close to that of *Fgf20*^-/-^ embryos (p>0.5). Cell death indexes were comparable to both *Fgf20*^-/-^ and *Fgf20*^-/-^;*Spry1*^-/+^ embryos (1.3±0.9, 1.0±0.4, n=4, respectively, ns) (Figs. 3O, P, U). Cell death index in *Fgf9*^-/+^;*Fgf20*^-/-^;*Spry1*^-/+^ embryos was increased compared to that of *Fgf9*^-/+^;*Fgf20*^-/-^ embryos (2.7±1.2, 1.1±0.3, n=4, respectively, p<0.05) (Figs. 3Q, R, U). Also, cell death index in *Fgf9*^-/-^;*Fgf20*^-/-^;Spry1^-/+^ embryos was increased compared to that of *Fgf9*^-/-^;*Fgf20*^-/-^ embryos (9.5±2.5, 2.6±1.8, n=4 respectively, p<0.001) (Figs. 3S, T, U). Together these data indicate that SPRY1 negatively regulates FGF9 and FGF20-dependent NPC survival and cell death.

**Fig. 3.**
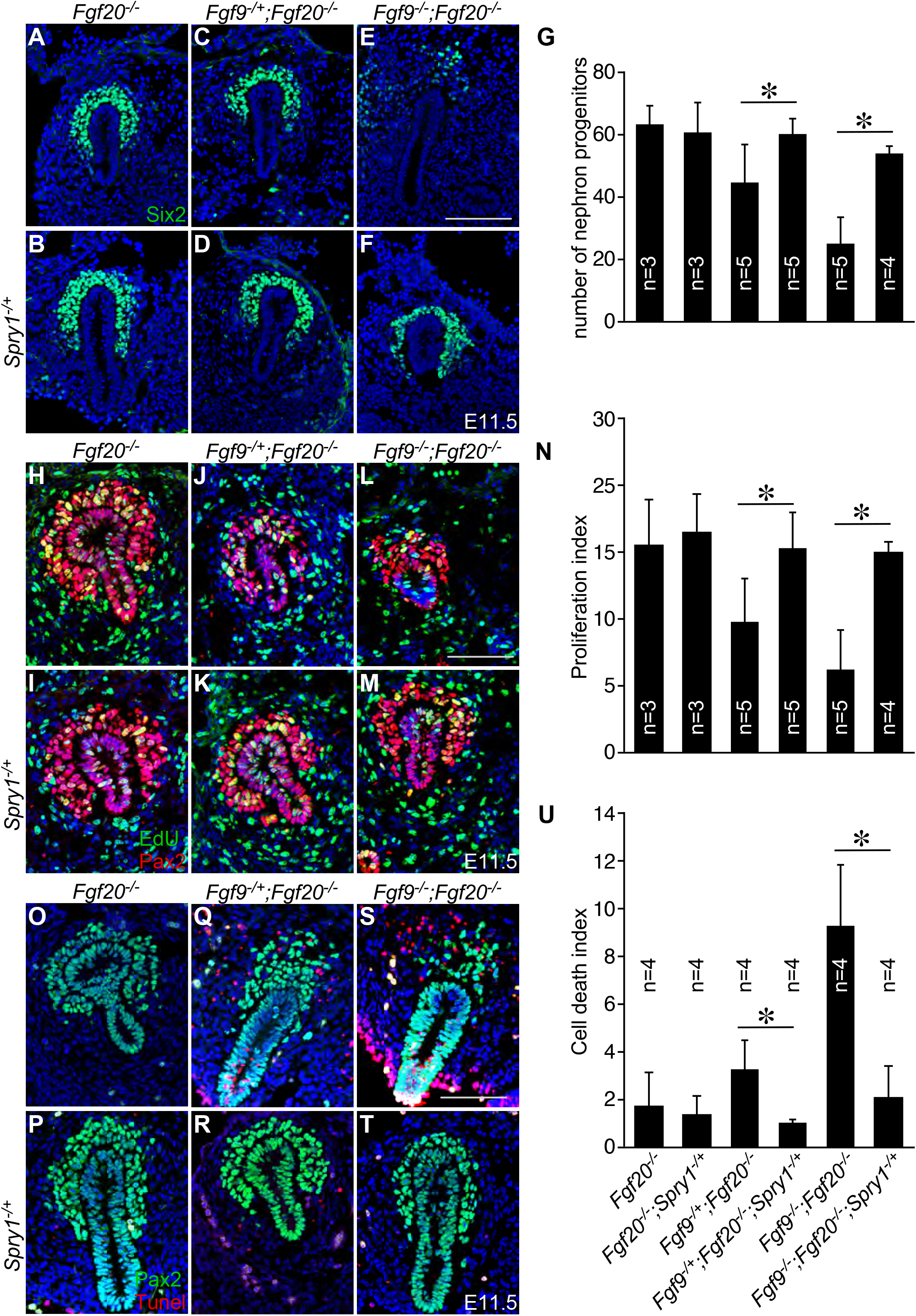
Haploinsufficiency of *Spry1* rescues nephron progenitor cell death and proliferation defects caused by loss of *Fgf9* and *Fgf20*. (**A-F**) Six2 staining of E11.5 *Fgf20*^-/-^ (**A**), *Fgf20*^-/-^;*Srpy1*^-/+^ (**B**), *Fgf9*^-/+^;*Fgf20*^-/-^ (**C**), *Fgf9*^-/+^;*Fgf20*^-/-^;*Spry1*^-/+^ (**D**), *Fgf9*^-/-^;*Fgf20*^-/-^ (**E**), and *Fgf9*^-/-^;*Fgf20*^-/-^;*Spry1*^-/+^ (**F**) kidney slides. (**G**) Number of nephron progenitors of *Fgf20*^-/-^ (n=3), *Fgf20*^-/-^;*Srpy1*^-/+^ (n=3), *Fgf9*^-/+^;*Fgf20*^-/-^ (n=5), *Fgf9*^-/+^;*Fgf20*^-/-^;*Spry1*^-/+^ (n=5), *Fgf9*^-/-^;*Fgf20*^-/-^ (n=5), and *Fgf9*^-/-^;*Fgf20*^-/-^;*Spry1*^-/+^ (n=4). (**H-M**) Pax2 and EdU staining of E11.5 *Fgf20*^-/-^ (**H**), *Fgf20*^-/-^;*Srpy1*^-/+^ (**I**), *Fgf9*^-/+^;*Fgf20*^-/-^ (**J**), *Fgf9*^-/+^;*Fgf20*^-/-^;*Spry1*^-/+^ (**K**), *Fgf9*^-/-^;*Fgf20*^-/-^ (**L**), and *Fgf9*^-/-^;*Fgf20*^-/-^;*Spry1*^-/+^ (**M**) kidney slides. (**N**) Proliferation index of *Fgf20*^-/-^ (n=3), *Fgf20*^-/-^;*Srpy1*^-/+^ (n=3), *Fgf9*^-/+^;*Fgf20*^-/-^ (n=5), *Fgf9*^-/+^;*Fgf20*^-/-^;*Spry1*^-/+^ (n=5), *Fgf9*^-/-^;*Fgf20*^-/-^ (n=5), and *Fgf9*^-/-^;*Fgf20*^-/-^;*Spry1*^-/+^ (n=4). (**O-T**) Pax2 and TUNEL staining of E11.5 *Fgf20*^-/-^ (**O**), *Fgf20*^-/-^;*Srpy1*^-/+^ (**P**), *Fgf9*^-/+^;*Fgf20*^-/-^ (**Q**), *Fgf9*^-/+^;*Fgf20*^-/-^;*Spry1*^-/+^ (**R**), *Fgf9*^-/-^;*Fgf20*^-/-^ (**S**), and *Fgf9*^-/-^;*Fgf20*^-/-^;*Spry1*^-/+^ (**T**) kidney slides. (**U**) Cell death index of *Fgf20*^-/-^ (n=4), *Fgf20*^-/-^;*Srpy1*^-/+^ (n=4), *Fgf9*^-/+^;*Fgf20*^-/-^ (n=4), *Fgf9*^-/+^;*Fgf20*^-/-^;*Spry1*^-/+^ (n=4), *Fgf9*^-/-^;*Fgf20*^-/-^ (n=4), and *Fgf9*^-/-^;*Fgf20*^-/-^;*Spry1*^-/+^ (n=4). Data is shown with mean±S.D. *P < 0.01. Scale bar, 100μm

### FGF8 functions together with FGF20 to regulate nephron progenitor maintenance

FGF8 is expressed in the NPCs and required for NPC survival and tubulogenesis at E14.5 (23–26). Since FGF8 is also required for NPC survival, we hypothesize that FGF8 functions together with FGF9 and FGF20 to maintain NPCs. To investigate this hypothesis, we deleted *Fgf8* in the NPC together with *Fgf20* using *Fgf20^Cre^* mouse line and analyzed kidneys. Kidneys in *Fgf8^fl/-^;Fgf20^Cre/+^* and *Fgf8^fl/+^;Fgf20^Cre/-^* embryos were smaller than those of *Fgf8^fl/+^;Fgf20^Cre/+^* embryos (Figs. 4A, B, C). *Fgf8^fl/-^;Fgf20^Cre/-^* embryos showed bilateral kidney agenesis (Fig. 4D). Kidneys of *Fgf8^fl/-^;Fgf20^Cre/+^* contained no nephrons which is consistent with previous publication (Fig. 4F) (23, 24). *Fgf8^fl/+^;Fgf20^Cre/-^* kidneys had less nephrons compared to *Fgf8^fl/+^;Fgf20^Cre/+^* embryos (Figs. 4E and G). UBs had less branching in both E14.5 *Fgf8^fl/-^;Fgf20^Cre/+^* and *Fgf8^fl/+^;Fgf20^Cre/-^* kidneys as notified by *Wnt9b* staining, the marker of UB (27) (Figs. S6A-C). *Wnt4* expression was diminished in the *Fgf8^fl/-^;Fgf20^Cre/+^* kidneys which is consistent with previous publication (Figs. 6E) (23, 24). However, *Wnt4* expression in the *Fgf8^fl/+^;Fgf20^Cre/-^* kidneys was comparable to *Fgf8^fl/+^;Fgf20^Cre/+^* (Figs. 6D and F) indicating that *Fgf20* does not regulate *Wnt4* expression.

**Fig. 4.**
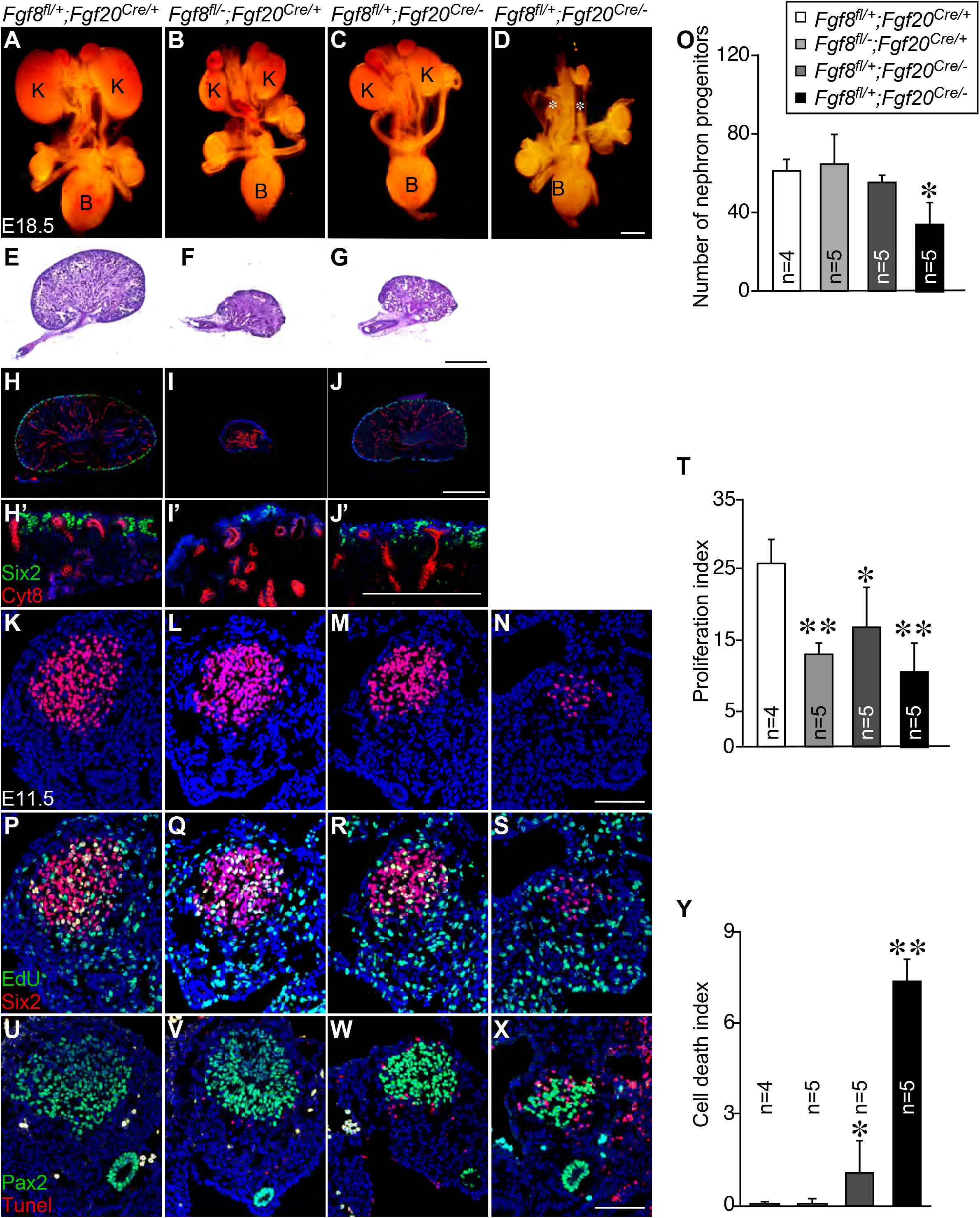
*Fgf8* functions redundantly with *Fgf20* to maintain nephrogenesis and nephron progenitors. (**A-D**) Morphology of urogenital system in E18.5 *Fgf8^fl/+^;Fgf20^Cre/+^* (**A**), *Fgf8^fl/-^;Fgf20^Cre/+^* (**B**), *Fgf8^fl/+^;Fgf20^Cre/-^* (**C**), and *Fgf8^fl/-^;Fgf20^Cre/-^* (**D**). (E-G) Hematoxylin & Eosin staining of *Fgf8^fl/+^;Fgf20^Cre/+^* (**E**), *Fgf8^fl/-^;Fgf20^Cre/+^* (**F**), and *Fgf8^fl/+^;Fgf20^Cre/-^* (**G**) kidneys. (**H-J**) Sections of E18.5 kidneys stained with Cytokeratin 8 (Cyt8) and Six2 in *Fgf8^fl/+^;Fgf20^Cre/+^* (**H**), *Fgf8^fl/-^;Fgf20^Cre/+^* (**I**), and *Fgf8^fl/+^;Fgf20^Cre/-^* (**J**) kidneys. (**H’-J’**) high power image of nephrogenic zone from *Fgf8^fl/+^;Fgf20^Cre/+^* (**H’**), *Fgf8^fl/-^;Fgf20^Cre/+^* (**I’**), and *Fgf8^fl/+^;Fgf20^Cre/-^* (**J’**) kidneys. (**K-N**) Six2 staining of E11.5 *Fgf8^fl/+^;Fgf20^Cre/+^* (**K**), *Fgf8^fl/-^;Fgf20^Cre/+^* (**L**), *Fgf8^fl/+^;Fgf20^Cre/-^* (**M**), and *Fgf8^fl/-^;Fgf20^Cre/-^* (**N**) kidney slides. (**O**) Number of nephron progenitors of *Fgf8^fl/+^;Fgf20^Cre/+^* (n=4), *Fgf8^fl/-^;Fgf20^Cre/+^* (n=5), *Fgf8^fl/+^;Fgf20^Cre/-^* (n=5), and *Fgf8^fl/-^;Fgf20^Cre/-^* (n=5). (**P-S**) Six2 and EdU staining of E11.5 *Fgf8^fl/+^;Fgf20^Cre/+^* (**P**), *Fgf8^fl/-^;Fgf20^Cre/+^* (**Q**), *Fgf8^fl/+^;Fgf20^Cre/-^* (**R**), and *Fgf8^fl/-^;Fgf20^Cre/-^* (**S**) kidney slides. (**T**) Proliferation index of *Fgf8^fl/+^;Fgf20^Cre/+^* (n=4), *Fgf8^fl/-^;Fgf20^Cre/+^* (n=5), *Fgf8^fl/+^;Fgf20^Cre/-^* (n=5), and *Fgf8^fl/-^;Fgf20^Cre/-^* (n=5). (**U-X**) Pax2 and TUNEL staining of E11.5 *Fgf8^fl/+^;Fgf20^Cre/+^* (**U**), *Fgf8^fl/-^;Fgf20^Cre/+^* (**V**), *Fgf8^fl/+^;Fgf20^Cre/-^* (**W**), and *Fgf8^fl/-^;Fgf20^Cre/-^* (**X**) kidney slides. (**U**) Cell death index of *Fgf8^fl/+^;Fgf20^Cre/+^* (n=4), *Fgf8^fl/-^;Fgf20^Cre/+^* (n=5), *Fgf8^fl/+^;Fgf20^Cre/-^* (n=5), and *Fgf8^fl/-^;Fgf20^Cre/-^* (n=5). K, kidney, B, bladder. * in D indicates loss of kidneys. Data is shown with mean±S.D. *P < 0.05. **P < 0.01. Scale bars in D, G, J, J’ 500μm. Scale bars in N, X, 100μm.

In NPC maintenance, Six2+ NPCs were decreased in E18.5 kidneys of *Fgf8^fl/-^;Fgf20^Cre/+^* and *Fgf8^fl/+^;Fgf20^Cre/-^* embryos compared to controls (Figs. 4H-J’). At E11.5, number of NPCs were comparable in *Fgf8^fl/+^;Fgf20^Cre/+^, Fgf8^fl/-^;Fgf20^Cre/+^, Fgf9^fl/+^;Fgf20^Cre/-^* embryos (61.8±5.5, 65.1±14.7, 55.9±3.0, respectively, n=4, 5, 5, respectively, ns)(Figs. 4K-M and O). However, number of NPCs in *Fgf8^fl/-^;Fgf20^Cre/-^* embryos (34.8±10.2, n=5, p<0.05) was significantly decreased compared to *Fgf8^fl/+^;Fgf20^Cre/+^* embryos (Figs. 4N and O). Proliferation rates of NPCs in *Fgf8^fl/-^;Fgf20^Cre/+^, Fgf8^fl/+^;Fgf20^Cre/-^*, and *Fgf8^fl/-^;Fgf20^Cre/-^* embryos (13.3±1.4, p<0.01, 17.0±5.5, p<0.05, 10.8±4.0, p<0.01, n=5, respectively) were significantly decreased compared to *Fgf9*^-/+^;*Fgf20*^-/+^ embryos (26.1±3.2, n=4) (Figs. 4P-S and T). Cell death rate of NPCs in *Fgf8^fl/-^;Fgf20^Cre/-^* embryos were comparable (0.1±0.1, ns) compared to controls (0.0±0.0) (Figs. 4U, V, and Y). Cell death rate of NPCs in *Fgf8^fl/+^;Fgf20^Cre^* embryos (1.1±1.0, n=5 p<0.05) embryos was increased (Figs. 4W and Y). Cell death rate of NPCs was further increased in *Fgf8^fl/-^;Fgf20^Cre/-^* (7.4±0.7, n=5, p<0.01) embryos compared to controls (p<0.01) and *Fgf8^fl/+^;Fgf20^Cre/-^* (p<0.01) embryos (Figs. 4X and Y). In addition, similar to *Fgf9*^-/-^;*Fgf20*^-/-^ (12), Pax2 expression in the NPCs were decreased in the *Fgf8^fl/-^;Fgf20^Cre/-^* embryos (Fig. S7A-D).

We also analyzed early UB branching. At E11, *Ret* was expressed in UBs and its expression was comparable to all the genotypes suggesting UB induction is not affected (Figs. S7E-H). At E11.5, *Ret+* UBs were bifurcated in *Fgf8^fl/+^;Fgf20^Cre/+^* embryo (Fig. S7J).

However, in *Fgf8^fl/-^;Fgf20^Cre/+^, Fgf8^fl/+^;Fgf20^Cre/-^*, and *Fgf8^fl/-^;Fgf20^Cre/-^* embryos, UBs were not yet bifurcated (Figs. S7J-L). *Gdnf* expression was comparable to *Fgf8^fl/+^;Fgf20^Cre/+^* and *Fgf8^fl/-^;Fgf20^Cre/+^* embryos (Figs. S7Q and R). However, expression of *Gdnf* was decreased in *Fgf8^fl/+^;Fgf20^Cre/-^* embryos (Fig. S7O) and diminished in *Fgf8^fl/-^;Fgf20^Cre/-^* embryos (Fig. S7P). In addition, *Etv4* and *Etv5* were diminished in *Fgf8^fl/-^;Fgf20^Cre/-^* embryos (Figs. S7Q-X). Together, these data indicate that *Fgf8*, similar to *Fgf9*, functions together with *Fgf20* to regulate kidney development.

### *Spry1* haploinsufficiency does not rescue kidney defects of *Fgf8* and *Fgf20* double knock-out

Next, we questioned whether loss of *Spry1* also rescue renal phenotypes caused by loss of *Fgf8* and *Fgf20*. Therefore, we generated *Spry1, Fgf8, Fgf20* compound mutants and analyzed kidneys. In *Fgf8^fl/-^;Fgf20^Cre/-^* embryos, among 16 kidneys, 2 very small kidneys were observed and 14 kidneys were missing (Figs. S8A and C). In *Fgf8^fl/-^;Fgf20^Cre/-^;*Spry1*^-/+^* embryos, 2 out of 10 kidneys were severely hypoplastic and 8 kidneys were missing (Figs. S8B and C). In addition, NPC number (34.8±10.2 in *Fgf8^fl/-^;Fgf20^Cre/-^*, n=5, and 35.4±1.7 in *Fgf8^fl/-^;Fgf20^Cre/-^;Spry1^-/+^*, n=3, ns), proliferation rate (10.8±4.0 in *Fgf8^fl/-^;Fgf20^Cre/-^*, n=5, and 10.3±2.3 in *Fgf8^fl/-^;Fgf20^Cre/-^;Spry1^-/+^*, n=3, ns), and cell death index (7.3±0.7 in *Fgf8^fl/-^;Fgf20^Cre/-^*, n=5, and 5.9±2.2 in *Fgf8^fl/-^;Fgf20^Cre/-^;Spry1^-/+^*, n=3, ns) were not rescued in the *Fgf8^fl/-^;Fgf20^Cre/-^;Spry1^-/+^* embryos compared to *Fgf8^fl/-^;Fgf20^Cre/-^* embryos (Figs. S8D-F).

### SPRY1 antagonizes FGFs in dose dependent manner

We identified that FGF8, FGF9, and FGF20 function together to regulate NPC maintenance. We also identified that deletion of *Spry1* did not rescue renal phenotypes caused by loss of both *Fgf8* and *Fgf20*. These results suggest that antagonism of SPRY1 to the FGF signal is limited by *Fgf* genes copy number. To investigate this, we generated *Fgf8, Fgf9, Fgf20*, and *Spry1* compound mutants and analyzed kidneys. In *Fgf8^fl/+^;Fgf9^-/+^;Fgf20^Cre/-^* embryos, 1 out of 10 kidney was normal, 6 were hypoplastic and 3 were missing (Figs. 5A and E). Kidney size was 34.8±34.3 (p<0.001) compared to *Fgf8^fl/+^;Fgf9^-/+^;Fgf20^Cre/-^Spry1^-/+^* embryos (Fig. 5F). In *Fgf8^fl/+^;Fgf9^-/+^;Fgf20^Cre/-^Spry1^-/+^* embryos, all kidneys (10 out of 10) were increased in size compared to *Fgf8^fl/+^;Fgf9^-/+^;Fgf20^Cre/-^* (Figs. 5B, E, F). All the kidneys (4 out of 4) in *Fgf8^fl/+^;*Fgf9*^-/-^;Fgf20^Cre/-^* embryos were missing (Figs. 5C, E, F). *Fgf8^fl/+^;*Fgf9*^-/-^;Fgf20^Cre/-^;*Spry1*^-/+^* embryos also lost all kidneys (Figs. 5C, D, E, F). These data indicate that rescue of kidney phenotypes in the haploinsufficiency of *Spry1* is depends on the dosage of gene copy numbers of *Fgfs*.

**Fig. 5.**
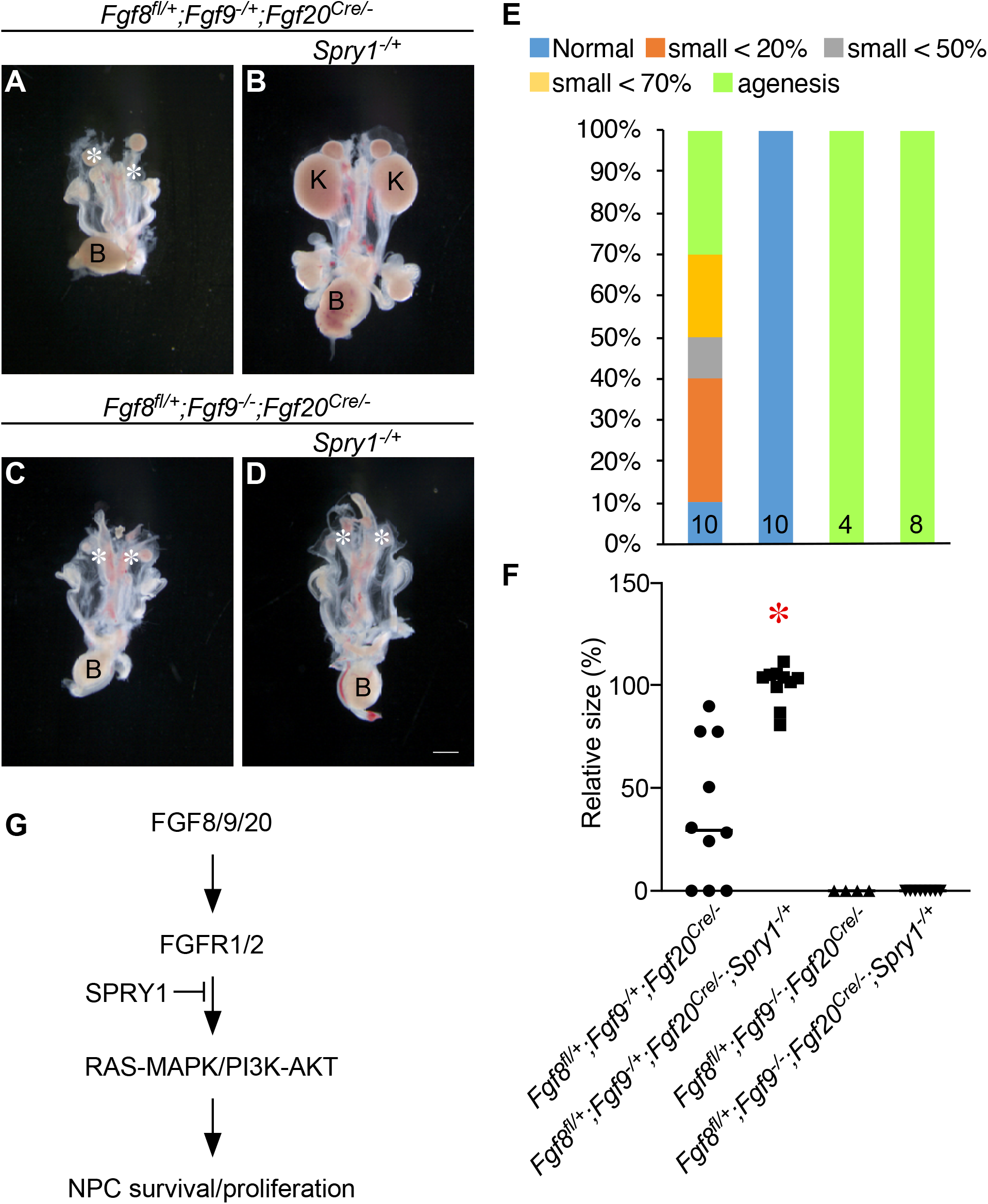
Haploinsufficiency of *Spry1* does not rescue renal phenotypes caused by loss of *Fgf8, Fgf9* and *Fgf20*. (**A-D**) Morphology of urogenital system in E18.5 *Fgf8^fl/+^;Fgf9^-/+^;Fgf20^Cre/-^* (**A**), *Fgf8^fl/+^;Fgf9^-/+^;Fgf20^Cre/-^*;*Spry1*^-/+^ (**B**), *Fgf8^fl/+^;Fgf9^-/-^;Fgf20^Cre/-^* (**C**), and *Fgf8^fl/+^;Fgf9^-/+^;Fgf20^Cre/-^;*Spry1*^-/+^* (**D**). (**E**) percentage of kidney phenotypes. (**F**) Relative kidney size of *Fgf8^fl/+^;Fgf9^-/+^;Fgf20^Cre/-^* (n=10), *Fgf8^fl/+^*;*Fgf9^-/+^*;*Fgf20^Cre/-^*;*Spry1*^-/+^ (n=10), *Fgf8^fl/+^;*Fgf9*^-/-^;Fgf20^Cre/-^* (n=4), and *Fgf8^fl/+^*;*Fgf9*^-/+^;*Fgf20^Cre/-^*;*Spry1*^-/+^ (n=8). (**G**) Model of FGFs and SPRY1 during NPC maintenance. K, kidney, B, bladder. * in A, C, and D indicate loss of kidneys. *P < 0.01. Scale bar: 100μm.

## Discussion

SPRY was originally identified to regulate development of *Drosophila* trachea through antagonizing branchless and breathless, which are orthologs of mammalian FGF and FGFR, respectively (28). The mammalian SPRYs antagonize FGF signaling in many developing organs including lung, mandible, external genitalia, long bone, auditory, and tooth (29–34). However, in kidney development, SPRY1 seems to function mainly as an antagonist of Ret-GDNF dependent UB branching. (15, 17). Of note, Brown et al, showed that ectopic expression of SPRY1 in NPCs resulted in loss of NPC due to NPC cell death (35), suggesting a possible role of SPRY1 in NPCs. Indeed, we present data indicating that haploinsufficiency of *Spry1* partially rescued renal phenotypes due to loss of *Fgf9* and *Fgf20* during NPC maintenance. NPC cell death, proliferation defect, and premature depletion was rescued. NPC specific *Spry1* deletion also rescued the *Fgf9* and *Fgf20* loss of function deletion indicating antagonistic function of *Spry1* is in a cell autonomous manner within NPCs.

FGF8 also functions together with FGF20 to maintain NPCs. At E11.5 *Fgf8* is expressed in the NPCs and its expression is restricted to pre-tubular aggregate (PTA) and renal vesicle as the embryo develops (23, 24). Previous studies indicate that *Fgf8* is required for NPC and PTA survival at later stage of development (E14.5 and later) but dispensable for NPC proliferation (23, 24). Interestingly, in this study, we identified that at early developmental stage (E11.5), proliferation of NPCs was decreased in all *Fgf8, Fgf20* compound mutants (*Fgf8^-/fl^;Fgf20^Cre/+^, Fgf8^-/+^;Fgf20^Cre/-^*, and *Fgf8^-/fl^;Fgf20^Cre/-^*). And NPC cell death was mostly pronounced in the *Fgf8* and *Fgf20* double mutant kidneys. *Fgf20* does not seem to function together with *Fgf8* to maintain PTA, as PTA marker (Wnt4) was not changed in the *Fgf8^fl/+^*;*Fgf20*^-/-^ kidneys. Different from *Fgf9* and *Fgf20*, deleting one copy of *Spry1* did not rescue renal phenotypes caused by loss of *Fgf8* and *Fgf20*. No kidneys developed in *Fgf8, Fgf20*, and *Spry1* compound mutant (*Fgf8^fl/-^;Fgf20^Cre/-^;Spry1^-/+^*). Also, no rescue of NPC proliferation and cell death occurs. More intriguingly, antagonism of SPRY1 to FGFs is dose dependent as haploinsufficiency of *Spry1* rescues renal agenesis in *Fgf8^fl/+^*;*Fgf9*^-/+^;*Fgf20*^-/-^ but not in the *Fgf8^fl/+^;*Fgf9*^-/-^;Fgf20^Cre/-^*. Based on this, we propose a model (Fig. 5G) of FGFs and SPRY1 functions for NPC maintenance during kidney development. FGF8, FGF9, and FGF20 bind to FGFR1 and FGFR2 in the NPCs to activate downstream signaling pathways, including RAS-MAPK and PI3K-AKT, that supports NPC proliferation, survival and stemness (35, 36). Upon activation of FGF signaling, *Spry1* is upregulated and fine-tune FGF signal by antagonizing RAS-MAPK and PI3K-AKT signaling cascade (37).

Another interesting observation is that *Gdnf, Etv4* and *Etv5* expression were lost in both *Fgf9/20* and *Fgf8/20* double mutants. As GDNF is critical activator of RET for UB branching morphogenesis and ETV4 and ETV5 are downstream effector of RET for UB branching (20), it would be interesting to investigate whether lowering FGF signal in both NPC and UB causes renal defects. FGF10 functions to promote UB branching in the absence of Ret-GDNF/SPRY1 (18). However, *Fgf10* null embryos has mild effect of UB branching generating small kidneys (38). Deleting both *Fgf10* and *Fgf20* would give an insight for the combination FGF function in both NPC and UB.

In conclusion, fine-tuning of FGF signaling is required for proper NPC maintenance and number of nephrons. In recent advent of kidney organoids, many protocols use FGF9 to promote NPC from intermediate mesoderm using both mouse and human embryonic stem cells (39–42). This information would provide importance of fine-tuning FGF signal during kidney development and may be used to make better organoid protocol.

## Materials and Methods

### Mice

*Spry1*^-/+^ (15), *Spry1^fl/+^* (MMRRC:029870) (15), *Fgf8^fl/+^* (43), *Fgf8*^-/+^ (23), *Fgf9*^-/+^ (44), *Fgf20*^-/+^ (45), *Fgf20^Cre/+^* (46), *Rosa^TdTomato/+^* (Jax#007905) were used. All the mice were maintained in the University of Nebraska Medical Center animal facility according to animal care regulations, and the Animal Care and Use Committee (protocol number 16-005-02-EP).

## Acknowledgements

We thank Dr. Gail Marin for the *Fgf8^fl^, Fgf8^-^*, and *Spry1*^-^ mice. This work was supported by the NIH P30 DK074038 (S.H). We are grateful to Dr. Kameswaran Surendran (Sanford Research) for helpful discussion.

## Supporting Materials and Methods

### Histology

For histological analysis, E18.5 kidneys were dissected from embryos, fixed overnight with 4% paraformaldehyde (PFA) overnight at 4°C, and dehydrated with ethanol gradients. Dehydrated kidneys were embedded in paraffin and sectioned. The paraffin embedded sections were de-paraffinized, hydrated and stained with hematoxylin and eosin. Stained slides were dehydrated, mounted, and photographed with Zeiss microscope.

### Immunohistochemistry

E10.5 and 11.5 embryos were incubated with 30% sucrose solution. For E18.5 and P0 samples, kidneys were isolated from embryos and incubated with series of sucrose solutions (10%, 20%, and 30%). Samples were frozen sectioned and stored at −80°C for storage. For antibody staining, sections were washed with PBST (PBS + 0.5% Triton-X 100) for 30 min at RT and incubated with blocking solution (5% donkey serum, in PBST) for 1hr at RT. Sections were incubated with primary antibodies in PBST with 1% donkey serum overnight at 4°C. Sections were washed 3X with PBS and incubated with secondary antibodies for 30min at RT. After washing 3X with PBS, slides were mounted with vectorshield (Vector labs). Images were acquired with Zeiss Axioimage Z2 equipped with ApoTome. Antibodies used were Six2 (Proteintech 1:500), Cytokeratin-8 (DSHB, 1:40), Biotylated-DBA (Vector labs 1:500) FoxD1 (Santa Cruz, 1:200), and Pax2 (Covance, 1:200). Secondary antibodies conjugated with Alexa488 and Alexa555 (Molecular Probes 1:500) were used.

### EdU and TUNEL staining

For EdU pulse-labeling, moms containing E10.5 and E11.5 were injected (xxx mg/mg). 2hrs after injection, moms were sacrificed and embryos were collected and processed for frozen section. Staining of EdU was followed by Click-iT® EdU imaging kit protocol (Invitrogen, C10338). TUNEL staining was performed according to the manufacturer’s recommendation (Roche, In situ Cell Death Dection Kit).

### In situ hybridization

For sectioning in situ hybridization, paraffinized slides were hydrated with Diethylpyrocarbonate (DEPC) treated ethanol series and water, washed with hybridization solution, and incubated overnight with digoxigenin-labeled *Spry1* probe. After washing, samples were incubated with anti-digoxigenin antibody conjugated with alkaline phosphatase (Roche) and the color reaction was performed using alkaline phosphate substrate (Roche). For whole mount in situ hybridization, embryos were dissected in DEPC treated PBS and fixed with 4% PFA. After washing, samples were dehydrated with methanol series and stored until usage. Probes used for these studies were *Wnt9b, Wnt4, Pax2, Ret, Gdnf, Etv4*, and *Etv5*.

### Statistical analysis

GraphPad Prism was used to perform a Welch’s t-test or a non-parametric Kruskal-Wallis test with Dunn’s test compensating for multiple comparisons where appropriate. A value of P < 0.05 was considered statistically significant. For comparative analysis of kidney sizes, kidney perimeter was measured and individual measurements were used for statistical analysis. Three or more animals from at least two independent experiments were examined.

## Supporting Figure Legends

**Fig. S1.**
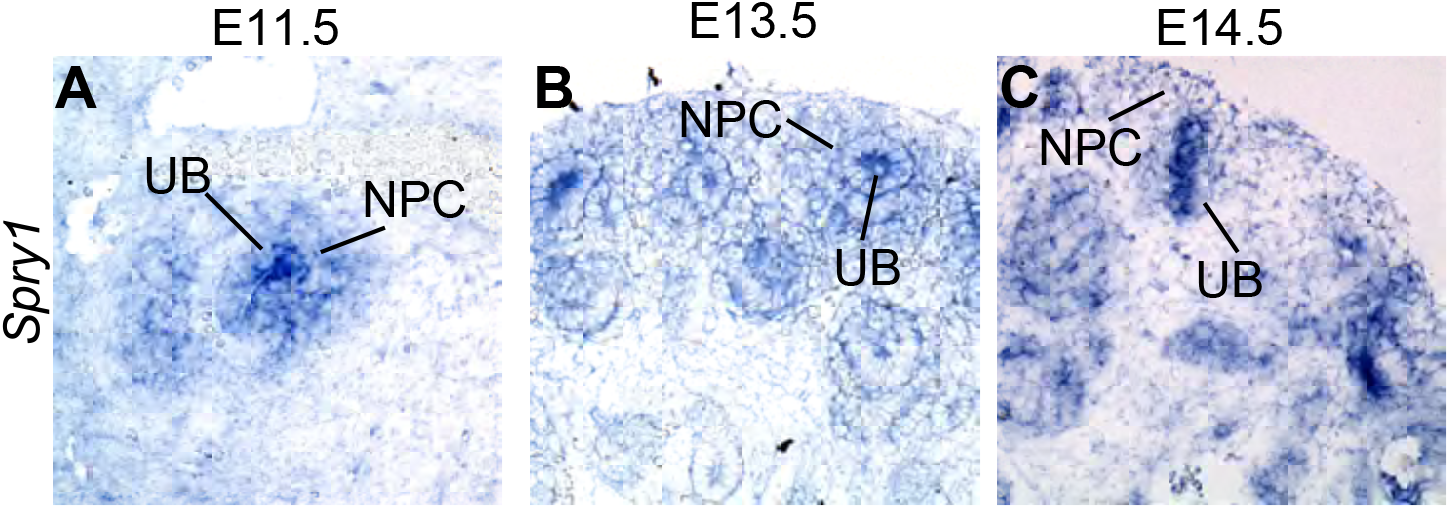
*In situ* hybridization of *Spry1*. (**A-C**) sectional *in situ* hybridization of *Spry1* in E11.5 (**A**), E13.5 (**B**), and E14.5 (**C**) kidneys. UB, ureteric bud. NPC, nephron progenitor cells.

**Fig. S2.**
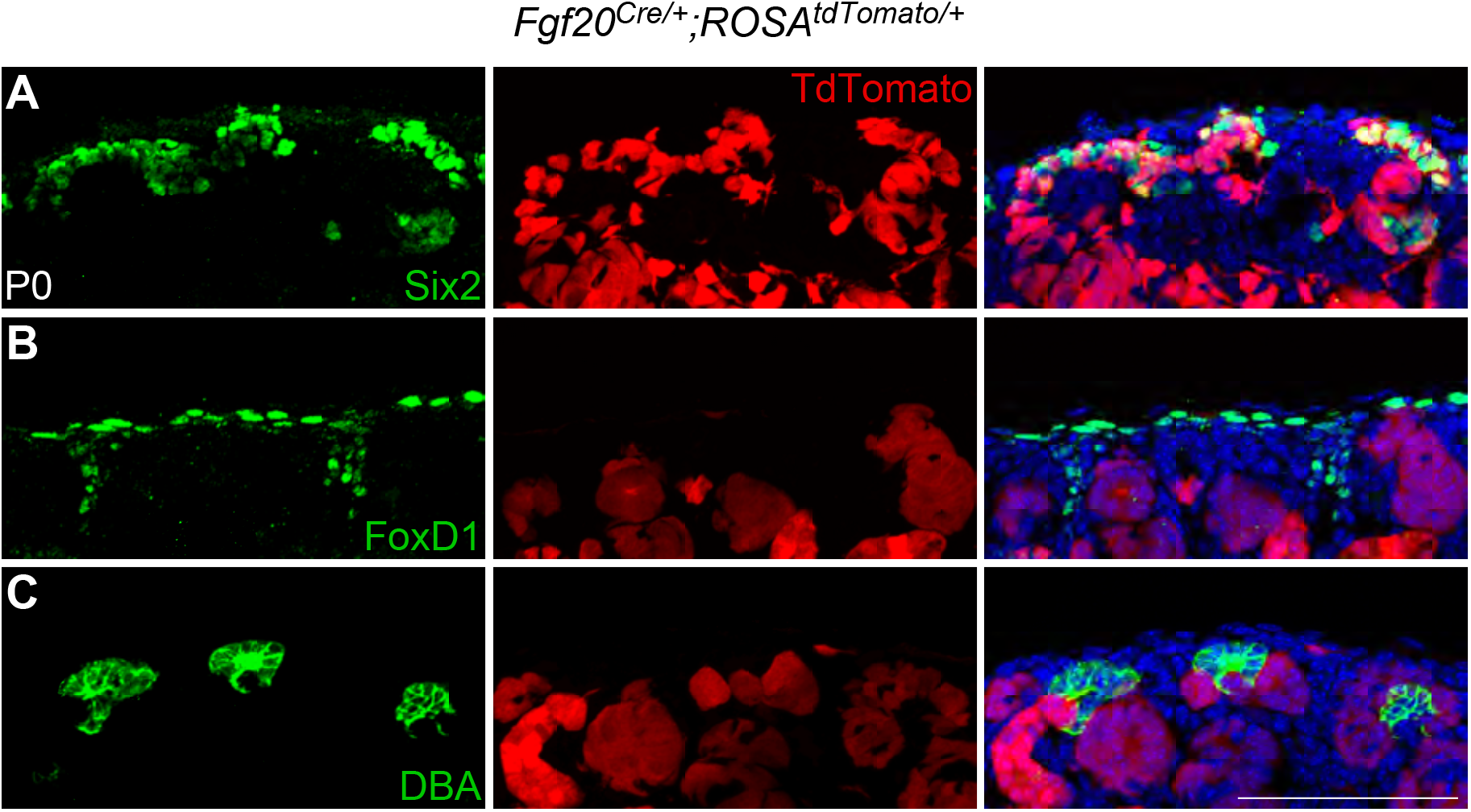
Lineage analysis of *Fgf20^Cre^*. (**A-C**) co-labeling Six2 (**A**), FoxD1 (**B**), and DBA (**C**) with tdTomato in P0 *Fgf20^Cre/+^;ROSA^tdTomato/+^* kidneys. Scale bar, 100μm.

**Fig. S3.**
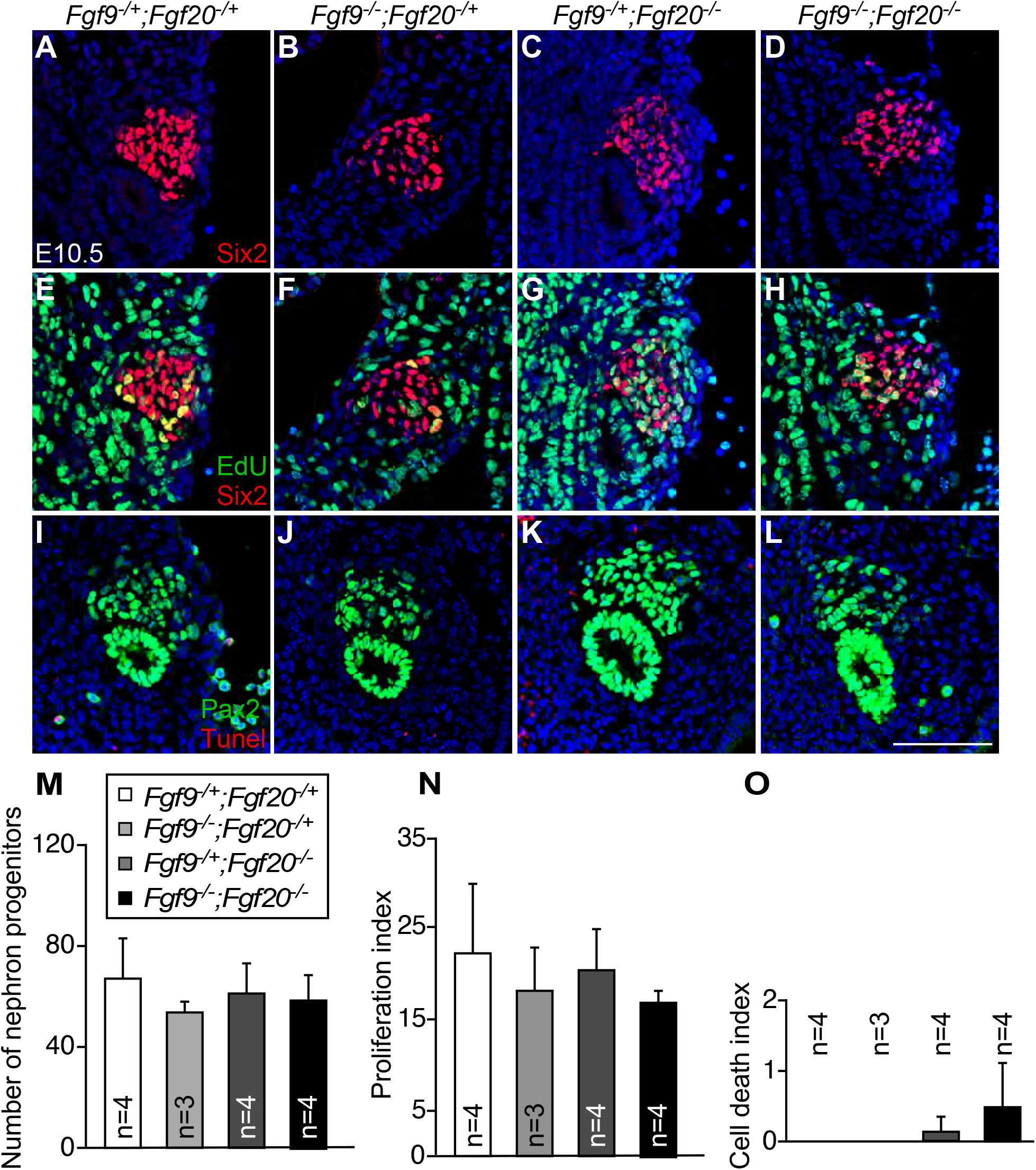
*Fgf9* and *Fgf20* are not required for cell death and proliferation of E10.5 nephron progenitors. (**A-D**) Six2 staining of E10.5 *Fgf9*^-/+^;*Fgf20*^-/+^ (**A**), *Fgf9*^-/-^;*Fgf20*^-/+^ (**B**), *Fgf9*^-/+^;*Fgf20*^-/-^ (**C**), and *Fgf9*^-/-^;*Fgf20*^-/-^ (**D**) kidney slides. (**E-H**) Six2 and EdU staining of E10.5 *Fgf9*^-/+^;*Fgf20*^-/+^ (**E**), *Fgf9*^-/-^;*Fgf20*^-/+^ (**F**), *Fgf9*^-/+^;*Fgf20*^-/-^ (**G**), and *Fgf9*^-/-^;*Fgf20*^-/-^ (**H**) kidney slides. (**I-L**) Pax2 and TUNEL staining of E10.5 *Fgf9*^-/+^;*Fgf20*^-/+^ (**I**), *Fgf9*^-/-^;*Fgf20*^-/+^ (**J**), *Fgf9*^-/+^;*Fgf20*^-/-^ (**K**), and *Fgf9*^-/-^;*Fgf20*^-/-^ (**L**) kidney slides. (**M**) Number of nephron progenitors of *Fgf9*^-/+^;*Fgf20*^/+^ (n=4), *Fgf9*^-/-^;*Fgf20*^-/+^ (n=3), *Fgf9*^-/+^;*Fgf20*^-/-^ (n=4), and *Fgf9*^-/-^;*Fgf20*^-/-^ (n=4). (**N**) Proliferation index of *Fgf9*^-/+^;*Fgf20*^-/+^ (n=4), *Fgf9*^-/-^;*Fgf20*^-/+^ (n=3), *Fgf9*^-/+^;*Fgf20*^-/-^ (n=4), and *Fgf9*^-/-^;*Fgf20*^-/-^ (n=4). (**O**) Cell death index of *Fgf9*^-/+^;*Fgf20*^-/+^ (n=4), *Fgf9*^-/-^;*Fgf20*^-/+^ (n=3), *Fgf9*^-/+^;*Fgf20*^-/-^ (n=4), and *Fgf9*^-/-^;*Fgf20*^-/-^ (n=4). Data is shown with mean±S.D. Scale bar, 100μm.

**Fig. S4.**
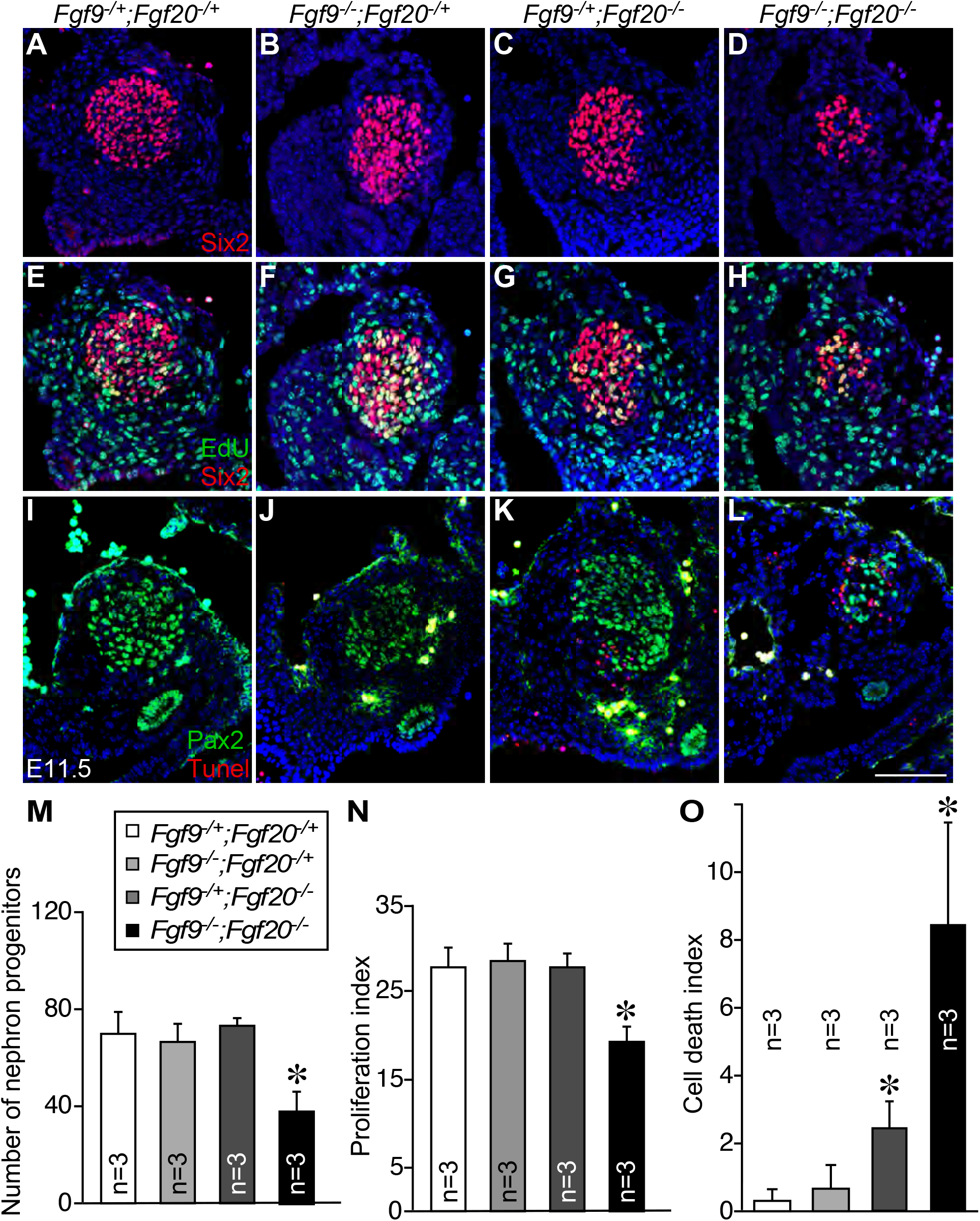
*Fgf9* and *Fgf20* are required for cell death and proliferation of E11.5 nephron progenitors. (**A-D**) Six2 staining of E11.5 *Fgf9*^-/+^;*Fgf20*^-/+^ (**A**), *Fgf9*^-/-^;*Fgf20*^-/+^ (**B**), *Fgf9*^-/+^;*Fgf20*^-/-^ (**C**), and *Fgf9*^-/-^;*Fgf20*^-/-^ (**D**) kidney slides. (**E-H**) Six2 and EdU staining of E10.5 *Fgf9*^-/+^;*Fgf20*^-/+^ (**E**), *Fgf9*^-/-^;*Fgf20*^-/+^ (**F**), *Fgf9*^-/+^;*Fgf20*^-/-^ (**G**), and *Fgf9*^-/-^;*Fgf20*^-/-^ (**H**) kidney slides. (**I-L**) Pax2 and TUNEL staining of E10.5 *Fgf9*^-/+^;*Fgf20*^-/+^ (**I**), *Fgf9*^-/-^;*Fgf20*^-/+^ (**J**), *Fgf9*^-/+^;*Fgf20*^-/-^ (**K**), and *Fgf9*^-/-^;*Fgf20*^-/-^ (**L**) kidney slides. (**M**) Number of nephron progenitors of *Fgf9*^-/+^;*Fgf20*^-/+^ (n=4), *Fgf9*^-/-^;*Fgf20*^-/+^ (n=3), *Fgf9*^-/+^;*Fgf20*^-/-^ (n=4), and *Fgf9*^-/-^;*Fgf20*^-/-^ (n=4). (**N**) Proliferation index of *Fgf9*^-/+^;*Fgf20*^-/+^ (n=4), *Fgf9*^-/-^;*Fgf20*^-/+^ (n=3), *Fgf9*^-/+^;*Fgf20*^-/-^ (n=4), and *Fgf9*^-/-^;*Fgf20*^-/-^ (n=4). (**O**) Cell death index of *Fgf9*^-/+^;*Fgf20*^-/+^ (n=4), *Fgf9*^-/-^;*Fgf20*^-/+^ (n=3), *Fgf9*^-/+^;*Fgf20*^-/-^ (n=4), and *Fgf9*^-/-^;*Fgf20*^-/-^ (n=4). Data is shown with mean±S.D. *P < 0.01. Scale bar, 100μm.

**Fig. S5.**
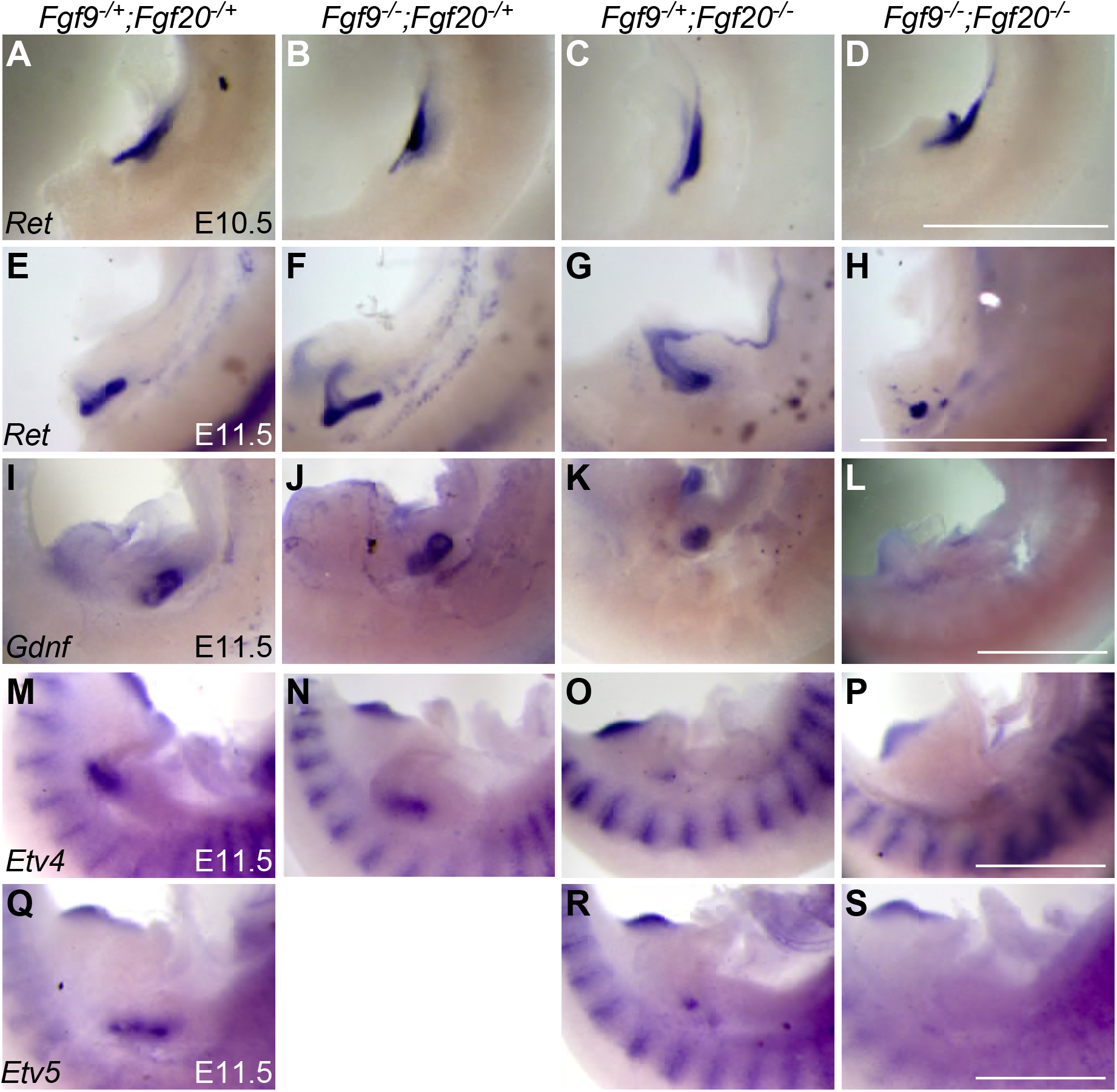
*Fgf9* and *Fgf20* regulate genes required for renal branching. (**A-H**) *Ret* in situ hybridization in E10.5 *Fgf9*^-/+^;*Fgf20*^-/+^ (**A**), *Fgf9*^-/-^;*Fgf20*^-/+^ (**B**), *Fgf9*^-/+^;*Fgf20*^-/-^ (**C**), and *Fgf9*^-/-^;*Fgf20*^-/-^ (**D**), and E11.5 *Fgf9*^-/+^;*Fgf20*^-/+^ (**E**), *Fgf9*^-/-^;*Fgf20*^-/+^ (**F**), *Fgf9*^-/+^;*Fgf20*^-/-^ (**G**), and *Fgf9*^-/-^;*Fgf20*^-/-^ (**H**) embryos. (**I-L**) *Gdnf* in situ hybridization of E11.5 *Fgf9*^-/+^;*Fgf20*^-/+^ (**I**), *Fgf9*^-/-^;*Fgf20*^-/+^ (**J**), *Fgf9*^-/+^;*Fgf20*^-/-^ (**K**), and *Fgf9*^-/-^;*Fgf20*^-/-^ (**L**) embryos. (**M-P**) *Etv4* in situ hybridization of E11.5 *Fgf9*^-/+^;*Fgf20*^-/+^ (**M**), *Fgf9*^-/-^;*Fgf20*^-/+^ (**N**), *Fgf9*^-/+^;*Fgf20*^-/-^ (**O**), and *Fgf9*^-/-^;*Fgf20*^-/-^ (**P**) embryos. (**Q-S**) *Etv5* in situ hybridization of E11.5 *Fgf9*^-/+^;*Fgf20*^-/+^ (**Q**), *Fgf9*^-/-^;*Fgf20*^-/+^ (**R**), and *Fgf9*^-/-^;*Fgf20*^-/-^ (**S**) embryos. Scale bar, 500μm.

**Fig. S6.**
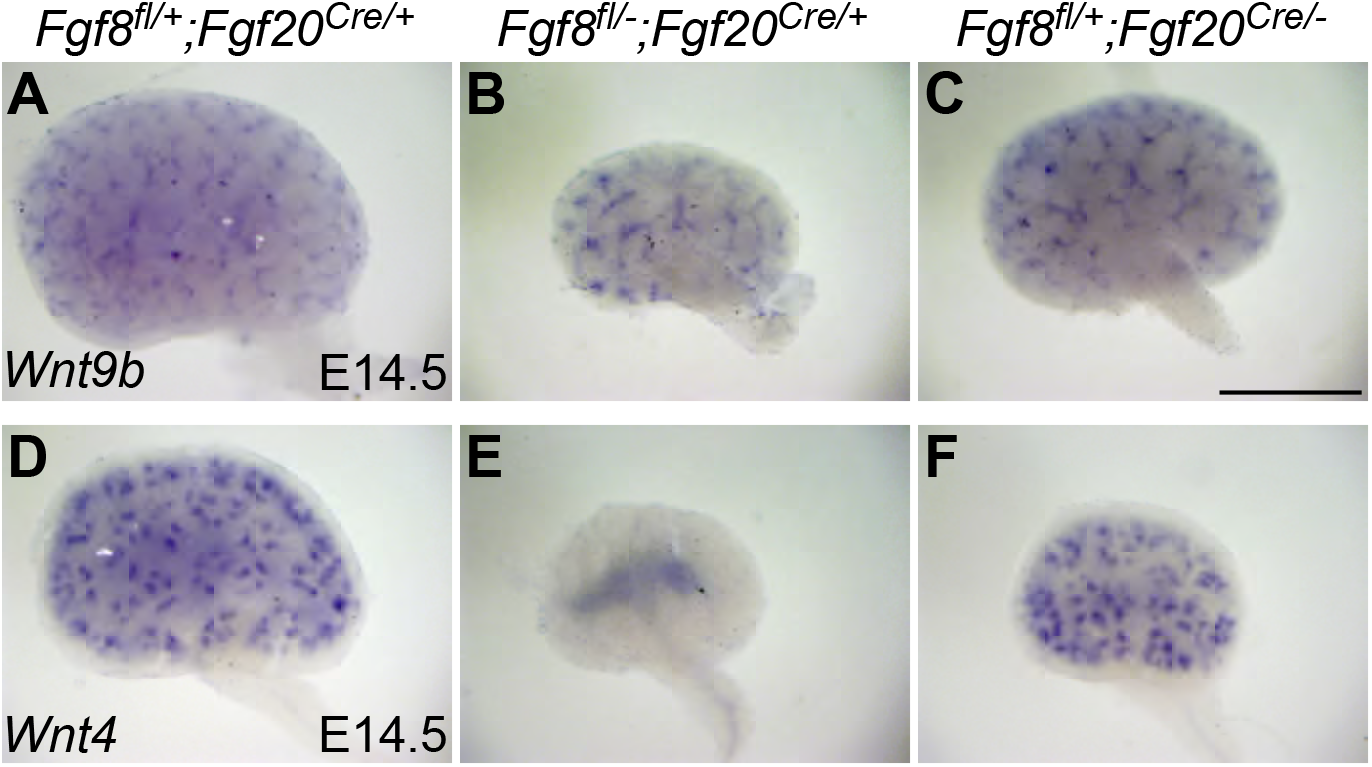
*Fgf20* does not affect renal phenotypes caused by loss of *Fgf8*. (**A-C**) *Wnt9b* in situ hybridization in E14.5 *Fgf8^fl/+^;Fgf20^Cre/+^* (**A**), *Fgf8^fl/-^;Fgf20^Cre/+^* (**B**), and *Fgf8^fl/+^;Fgf20^Cre/-^* (**C**) kidneys. (**D-F**) *Wnt4* in situ hybridization in E14.5 *Fgf8^fl/+^;Fgf20^Cre/+^* (**D**), *Fgf8^fl/-^;Fgf20^Cre/+^* (**E**), and *Fgf8^fl/+^;Fgf20^Cre/-^* (**F**) kidneys. Scale bar, 500μm.

**Fig. S7.**
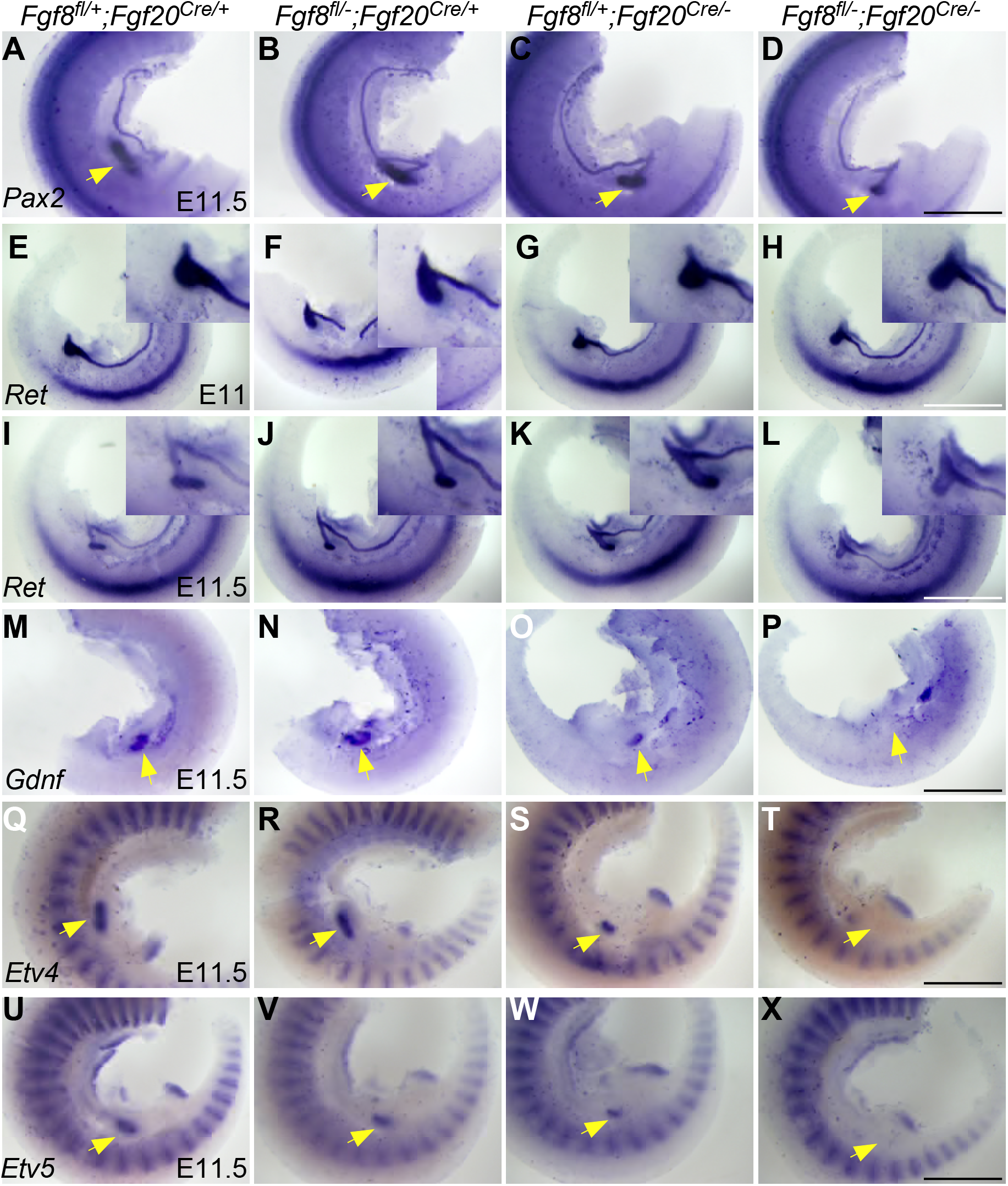
*Fgf8* and *Fgf20* regulate genes required for renal branching. (**A-D**) *Pax2* in situ hybridization in E11.5 *Fgf8^fl/+^;Fgf20^Cre/+^* (**A**), *Fgf8^fl/-^;Fgf20^Cre/+^* (**B**), *Fgf8^fl/+^;Fgf20^Cre/-^* (**C**), and *Fgf8^fl/-^;Fgf20^Cre/-^* (**D**) embryos. (**E-L**) *Ret* in situ hybridization in E11 *Fgf8^fl/+^;Fgf20^Cre/+^* (**E**), *Fgf8^fl/-^;Fgf20^Cre/+^* (**F**), *Fgf8^fl/+^;Fgf20^Cre/-^* (**G**), and *Fgf8^fl/-^;Fgf20^Cre/-^* (**H**), and E11.5 *Fgf8^fl/+^;Fgf20^Cre/+^* (**I**), *Fgf8^fl/-^;Fgf20^Cre/+^* (**J**), *Fgf8^fl/+^;Fgf20^Cre/-^* (**K**), and *Fgf8^fl/-^;Fgf20^Cre/-^* (**L**) embryos. (**M-P**) *Gdnf* in situ hybridization of E11.5 *Fgf8^fl/+^;Fgf20^Cre/+^* (**M**), *Fgf8^fl/-^;Fgf20^Cre/+^* (**N**), *Fgf8^fl/+^;Fgf20^Cre/-^* (**O**), and *Fgf8^fl/-^;Fgf20^Cre/-^* (**P**) embryos. (**Q-T**) *Etv4* in situ hybridization of E11.5 *Fgf8^fl/+^;Fgf20^Cre/+^* (**Q**), *Fgf8^fl/-^;Fgf20^Cre/+^* (**R**), *Fgf8^fl/+^;Fgf20^Cre/-^* (**S**), and *Fgf8^fl/-^;Fgf20^Cre/-^* (**T**) embryos. (**U-X**) *Etv5* in situ hybridization of E11.5 *Fgf8^fl/+^;Fgf20^Cre/+^* (**U**), *Fgf8^fl/-^;Fgf20^Cre/+^* (**V**), *Fgf8^fl/+^;Fgf20^Cre/-^* (**W**), and *Fgf8^fl/-^;Fgf20^Cre/-^* (**X**) embryos. Scale bar, 500μm.

**Fig. S8.**
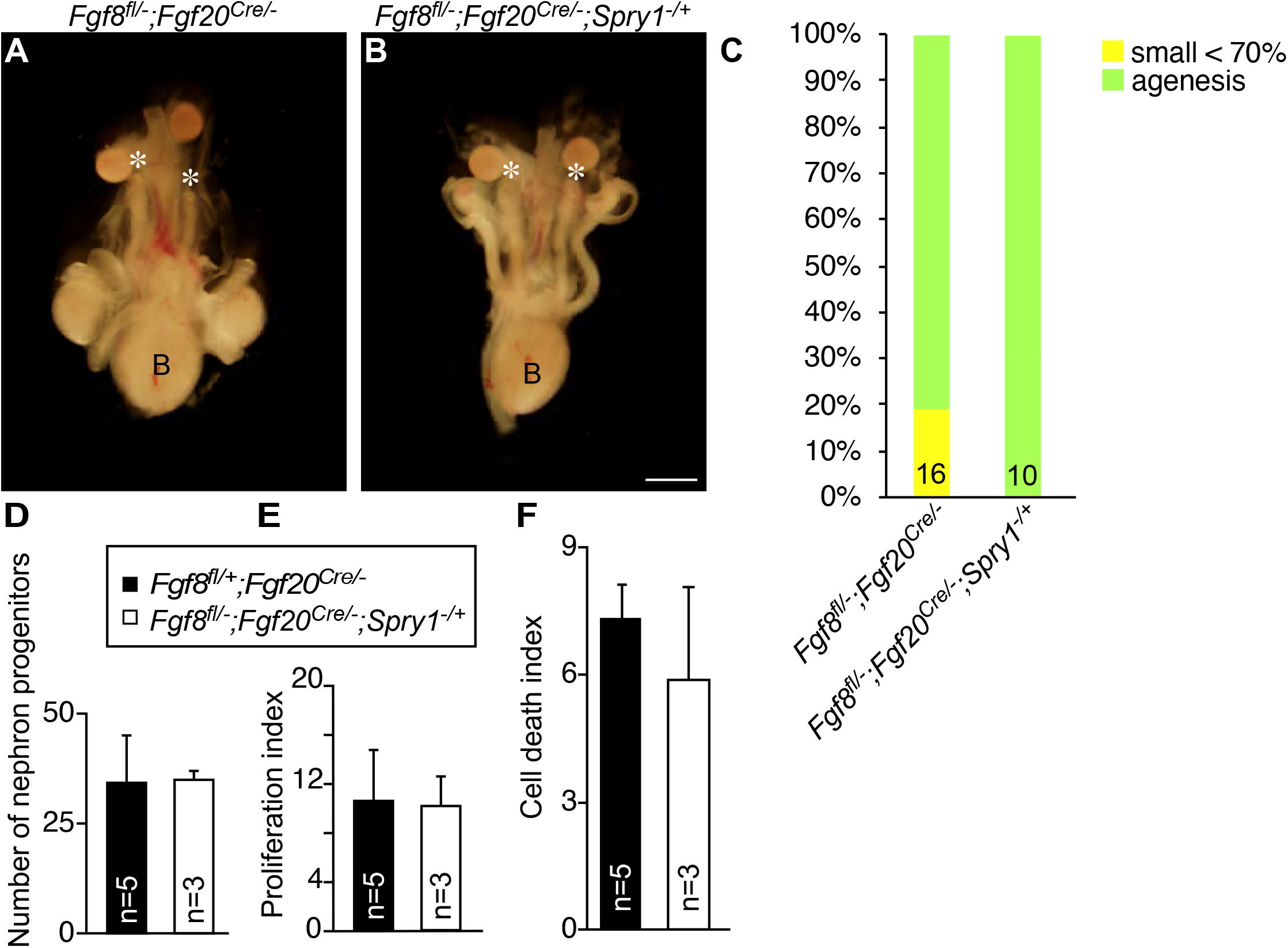
Haploinsufficiency of *Spry1* does not rescue renal phenotypes caused by loss of *Fgf8* and *Fgf20*. Morphology of urogenital system in E18.5 *Fgf8^fl/-^;Fgf20^Cre/-^* (**A**) and *Fgf8^fl/-^;Fgf20^Cre/-^*;*Spry1*^-/+^ (**B**). (**C**) percentage of kidney phenotypes. (**D**) Number of nephron progenitors of *Fgf8^fl/-^;Fgf20^Cre/-^* (n=5) and *Fgf8^fl/-^;Fgf20^Cre/-^*;*Spry1*^-/+^ (n=3). (**E**) Proliferation index of *Fgf8^fl/-^;Fgf20^Cre/-^* (n=5) and *Fgf8^fl/-^;Fgf20^Cre/-^*;*Spry1*^-/+^ (n=3). (**F**) Cell death index of *Fgf8^fl/-^;Fgf20^Cre/-^* (n=5) and *Fgf8^fl/-^;Fgf20^Cre/-^*;*Spry1*^-/+^ (n=3). B, bladder. * in A and B indicate loss of kidneys. Data is shown with mean±S.D. *P < 0.01. Scale bar, 100μm.

